# PKD2 structural destabilization drives primary cilia degeneration and ADPKD pathogenicity

**DOI:** 10.64898/2026.06.24.734313

**Authors:** Patricia Outeda, Qinzhe Wang, Thuy Vien, Orhi Esarte Palomero, Louise F. Kimura, Perry Summers, Terry J. Watnick, Feng Qian, Erhu Cao, Paul G. DeCaen

## Abstract

Human variants in renal polycystins (PKD1, PKD2) are responsible for most forms of autosomal dominant polycystic kidney disease (ADPKD), a common genetic disorder without curative drug treatment. Renal polycystins form ion channels in primary cilia, but our understanding of their molecular dysregulation caused by disease-associated variants is limited. Using cryo-electron microscopy (cryo-EM), primary cilia electrophysiology and super-resolution analysis, we investigated the mechanistic impact and pathogenic potential of the disease-associated PKD2 missense variant (D511V) located within the channel’s voltage sensor domain (VSD). Our findings define how this mutation neutralizes critical transmembrane charge interactions, which attenuates PKD2 protein stability resulting in abolished ciliary channel trafficking and function in membranes. To assess the pathogenic effect of this variant *in vivo*, we generated novel mouse strains carrying the analogous PKD2 mutation in combination with a conditional floxed allele (*Pkd2^D509V/fl^*) that exhibit renal tubule primary cilia degeneration and develop rapid renal cysts. Our results establish a clear direct correlation between the *in vitro* molecular dysfunction and phenotypic *in vivo* consequences while providing a valuable tool to evaluate ADPKD therapeutic interventions.

**Translational Statement:** ADPKD is a genetic kidney disorder affecting millions of patients globally and is primarily caused by variants in renal polycystin genes (PKD1, PKD2). Polycystins function as ion channel subunits in primary cilia but the mechanistic impact and cystogenic propensity of disease-associated variants remain poorly defined. The authors employ advanced methodologies including cryo-EM to uncover distinct structurally destabilizing effects of a human PKD2 mutation, while generating a new mouse model which genetically expresses the same variant and recapitulates the human disease. The findings define primary cilia degeneration results from PKD2 hypostasis and establish new tools to assess ADPKD therapies.

**Graphical Abstract:** 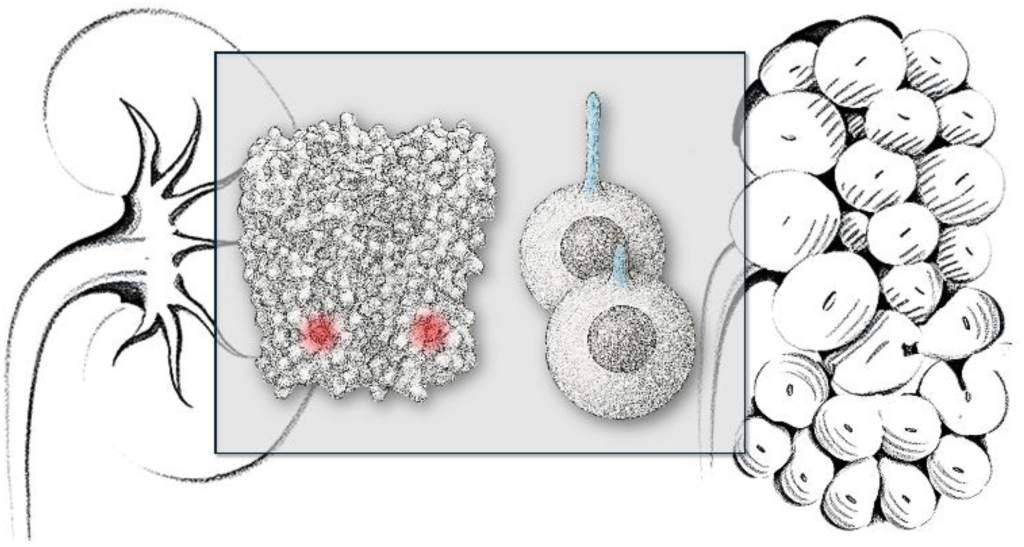

## INTRODUCTION

#### BOX 1. The polycystin nomenclature

The revised and current IUPHAR/BPS nomenclature creates ambiguity regarding the genetic identity of the polycystin family members of transient receptor potential ion channels, especially when cross-referencing manuscripts that describe subunits using the former system^1^. Traditionally, the products of polycystin genes (e.g., PKD2) are referred to as polycystin proteins (e.g., polycystin-2). For simplicity and to prevent confusion, we will refer to the polycystin gene name rather than differentiating gene and protein with separate names–a nomenclature we have recently outlined^2^. For differentiating between human variants and mouse models expressing analogous polycystin gene mutations, we will use the established convention of capital (e.g., PKD2) and lowercase italic (e.g., *Pkd2*) nomenclature, respectively.

Autosomal dominant polycystic kidney disease (ADPKD) is a prevalent monogenetic disorder characterized by progressive development of nephrotic cysts that ultimately culminate in fatal kidney failure^3^. Affecting over 12 million people globally, ADPKD currently has no curative drug treatments. The majority of ADPKD cases are attributed to genetic mutations in the renal polycystin ion channel subunits, PKD1 and PKD2. Approximately 80% of patients harbor germline variants in the PKD1 gene, while the remaining 15% carry mutations in *PKD2*^4,5^. Evidence from patient studies and animal models indicates that cyst formation is triggered by a second somatic mutation (“second hit”) in the remaining functional allele of either PKD1 or PKD2^6,7^. This genetic mechanism explains the progressive accumulation of focal renal cysts and the typical onset of kidney failure in midlife^5,8^.

While previous studies have explored the relationship between genotype and renal function in a limited subset of ADPKD patients, our understanding of the specific cystogenic potential of different types of mutations (e.g., missense, nonsense) and their genomic loci remains speculative^9^. Beyond the kidneys, polycystins play crucial roles in the development and maintenance of essential organ systems, including the cardiovascular system. In mice, bi-allelic loss of PKD1 or PKD2 expression causes embryonic lethality due to severe lymphatic, cardiovascular, hepatic and renal defects^10–14^. Mouse models with hypomorphic expression or conditional loss of polycystin alleles have faithfully reproduced ADPKD phenotypes offering valuable insights into disease mechanisms^15–20^. Although murine models harboring human disease-associated PKD1 variants have been developed^21^, the study of equivalent PKD2 mutations are relatively underexplored^22^. This study seeks to address this gap by investigating the molecular pathology and cystogenic potential of a specific PKD2 variant (D511V) identified in the ADPKD population. Through this research, we hope to enhance our understanding of the mutation-specific contributions to disease progression and its systemic impact.

Renal polycystins form homomeric (PKD2) and heteromeric (PKD1-PKD2) channel complexes, each with apparently distinct gating properties in primary cilia membranes^23–26^. Primary cilia reside on the apical surface of renal epithelial principal cells, where they coordinate signaling pathways that maintain tubular architecture, epithelial polarity, and nephron integrity^27,28^. While the characteristics of native heteromeric PKD1-PKD2 channels are poorly understood, recent studies suggests that these complexes might be proteolytically activated, but these effects are only observed after introducing non-native cleavage and gain-of-function mutations^29^. In contrast, homomeric PKD2 channels are voltage-gated, internal Ca^2+^ modulated and are readily measured from primary cilia membranes using submicron diameter electrodes, and when reconstituted from cellular membrane fractions, or when synthetically derived using cell-free systems^23,24,30^. Given their tenability to these methodologies, we will primarily consider the impact of PKD2 variants within the context of homomeric channels.

High resolution cryo-EM structural determination of homomeric PKD2 channels has provided a molecular context for interpreting the dysfunctional mechanistic impact of ADPKD-causing variants^25,31–33^ (**Supplemental Figure 1A**). Each of the four PKD2 subunits containing six transmembrane segments (S1-S6) that can comprise functional domains of the pore (PD), voltage sensor (VSD), tetrameric oligomerization domain of polycystins (TOP), and C-terminus (CTD). Most pathogenic missense variants (non-truncating) impact the TOP domain, which destabilizes this external region of the channel and results in gating defects, whereas those affecting the VSD (L517R, D511V) produce completely non-functional channels when reconstituted in bilayers (**Supplemental Figure 1B**).^18,22,34–36^. However, the underlying cause of the “channel-dead” properties and the variant capacity to induce cyst formation is largely unexplored. In this manuscript, we endeavor to discover the mechanistic impact of the D511V missense variant by examining its effects on channel expression, structural stability, trafficking, function and primary cilia maintenance using *in vitro* and *in vivo* methods. Our developed mouse strains harbor the analogous PKD2 mutation and reveal new insights into the molecular mechanism of polycystin dysfunction while providing a cystogenic model which mimics human ADPKD genetics and disease progression.

## METHODS

### Expression and purification of human PKD2 for D511V cryo-EM structural determination

The human PKD2 WT and D511V mutant constructs, encompassing residues 53–792, was expressed with an N-terminal maltose binding protein (MBP) fusion in HEK293S GnTI− (ATCC CRL-3022) cells using the BacMam system as described^37^. In brief, HEK293S GnTI−/− cells, grown in suspension in freestyle 293 expression medium (Invitrogen, Carlsbad, CA) at 37 °C in an orbital shaker, were transduced with PKD2 baculoviruses when cell density reached ∼2 × 10^6^/ml. Eight to twenty-four hours post transduction, sodium butyrate was added to the culture to a final concentration of 5 mM to enhance protein expression; temperature was reduced to 30 °C. Cells were harvested 72 h post transduction for subsequent affinity purification with amylose resin (New England BioLabs, Ipswich, MA). Human PKD2 protein was eluted from amylose resin with buffer composed of (in mM) 20 HEPES (pH 7.4), 150 NaCl, 0.5 TCEP, 0.5 MNG-3, 0.1 CHS, 20 maltose, 0.1 mg/ml soybean lipids, 10% glycerol. MBP tag was removed by TEV protease cleavage overnight at 4 °C. PKD2 was separated with a Superose 6 column using buffer composed of (in mM) 20 HEPES (pH 7.4), 150 NaCl, 0.5 TCEP, 25 × 10−3 MNG-3, and 5 × 10−3 CHS, and peaks corresponding to PKD2 homotetramer were pooled and concentrated for cryo-EM analyses.

### Cryo-EM data acquisition, image processing 3D reconstruction and model building

Each 3 μL of PKD2 sample at ∼1.5 mg/mL was applied to glow-discharged UltrAuFoil 1.2/1.3 holey 300-mesh gold grids. Grids were plunge-frozen in liquid ethane using a Vitrobot Mark III (FEI) set to 4 °C, 85% relative humidity, 20 s wait time, −1-mm offset, and 2.5 s blotting time. Cryo-EM datasets were collected on a 300-kV Titan Krios electron microscope (Thermo Fisher Scientific) equipped with a K2 Summit detector (Gatan) at a nominal magnification of 22,500x in super-resolution counting mode. Automatic data collection was performed using SerialEM^38^, with a defocus range between −0.6 and −3.6 μm. Movie frames were aligned, dose-weighted, and then summed into a single micrograph using MotionCor2^39^. Contrast transfer function parameters for micrographs were determined using the program CTFFIND4^40^. An estimated 1,500 particles were manually boxed out in RELION to generate initial 2D averages as a sanity check to confirm the presence of different tetrameric channel views. The well centered particles were then used to train an AI based autopicker (TOPAZ) to automatically pick particles from all micrographs^41^. All 2D and 3D classifications and refinements were performed in Cryosparc. Obvious “junk” particles (eg. ice contamination and gold support) were excluded from downstream processing based on results from one round of 2D classification. For the PKD2 D511V dataset, a total of 2,742,864 particles were extracted and multiple rounds of 2D classification in cryoSPARC v3 software^42^ removed most broken D511V particles or large aggregates, resulting in 398,779 good particles for multi-class *ab initial* model generation in Cryosparc. Then the same particles were used for a 6-class heterogenous refinement in C1 symmetry. The subset of particles from good 3D classes were combined for non-uniform refinement with C4 symmetry, resulting in a 3.1 Å reconstruction (local resolution ranges from 2.67 Å to 11.5 Å). For model building, the D511V map was locally sharpened in PHENIX1.20^43^ to enhance the side-chain density. Initial PKD2 wild type model (PDB: 5T4D) was manually docked into the EM density. 511D were mutated to 511V and missing loops were manually built in Coot^44^. Surprisingly our D511V map has good density for multiple N-glycans, so extended glycan structures were built with the N-glycan module^45^. The model was refined in real space against the locally sharpened maps using PHENIX 1.20 and assessed in MolProbity as shown in **Supplemental Table 1**. Figures were generated using UCSF ChimeraX^46^.

### Thermal Stability Assay

The fluorescence detection size-exclusion based thermostability assay was performed as previously described with minor modifications^47^. Approximately 9 μg of purified human PKD2 channel proteins (WT or D511V) in 30 μL FSEC buffer composed of 20 mM Hepes, 150 mM NaCl, 2 mM CaCl2, 0.5 mM TCEP, and 0.5 mM DDM, at pH 7.4, were incubated at 30 °C to 97.4 °C (30 °C, 32 °C, 35.2 °C, 39.3 °C, 44.9 °C, 49 °C, 50 °C, 51.9 °C, 52.1 °C, 54 °C, 55.4 °C, 59.4 °C, 64.9 °C, 69.2 °C, 72.1 °C, 74 °C, 75 °C, 77.4 °C, 80.6 °C, 84.5 °C, 90 °C, 94.4 °C, 97.4 °C) for 10 min in a thermal cycler. The treated samples were flash frozen in liquid nitrogen and thawed on ice to induce aggregation. After 10-fold sample dilution in the same buffer, the treated samples were cleared by ultracentrifugation at 40,000 rpm for 30 min. Thirty microliters of each cleared channel samples were separated on an analytical size-exclusion column (Superose 6 5/150 GL; GE Healthcare) at 0.3 mL/min flow rate. Proteins were detected by tryptophan fluorescence. The fluorescence-detection size-exclusion chromatography-based thermostability experiments were performed in triplicates for each temperature point. The peak height of the channel tetramer peak at different temperature points is normalized to that at lowest temperature to generate the thermal stability plot.

### Generation of HEK PKD1^Null^:PKD2^Null^ cells lines and the expression of PKD2 variants

Using our previously generated CRISPR/Cas9 gene edited HEK PKD2^null^ cell line, we introduced nonsense mutations to both PKD1 alleles to create a PKD1^null^:PKD2^null^ cell line. HEK 293 PKD2^null^ cells were electro-transfected with PKD1 sgRNAs (caccGCATAGGTGTGGTTGGCAGC and aaacGCTGCCAACCACACCTATGC) with the All-in-one Cas9 plasmid (Addgene)^48,49^. Cells generated from single cell clones were selected after 4 weeks of expansion under puromycin selection in a 96-wells plate. A HEK PKD1^Null^:PKD2^Null^ clone was verified by extracting the genomic DNA and sequencing for the introduced STOP codons within PKD1 and PKD2 genes using the following primers: PKD1 fwd (CTGATGGCTTAGGCCCCTACTG); PKD1 rev (CCTGGGTCTCGGTAGATGAACG); PKD2 fwd (AGCCTCAGGGCACAGAACAG); and PKD2 rev (CCACACTGCCCTTCATTGGC). HEK PKD1^Null^:PKD2^Null^ clones stably expressing the ARL13B-GFP cilia reporter (VectorBuilder, pLV[Exp]-Puro-CMV) were established after (2 μg/ml) for 30 to 90 days or 9-10 rounds of puromycin culture selection. Cells were then fluorescence-activated cell sorted (BD FacsMelody) at 5000 to 10,000 counts per minute to enrich for the transgene expression. To generate the WT and variant PKD2 N-terminally HA-tagged variant cell lines, the human PKD2 gene was subcloned into mammalian expression vector pRP[Exp]-CMV>HA (VectorBuilder). Missense variants were generated using standard site-directed mutagenesis. Cell lines were cultured in Dulbecco’s modified essential medium (DMEM) supplemented with 10% fetal bovine serum (FBS) and 100 units/ml penicillin, and 100 units/ml streptomycin and/or 1 μg/ml puromycin selection antibiotic.

### Electrophysiology recordings of PKD2 from primary cilia membranes

The electrophysiologist was blinded by a third party in the laboratory, whereby test groups were assigned a letter to conceal the genetic identity of the cells being evaluated. The identity of the cells remained unknown by the electrophysiologist until the analysis was complete. Ciliary ion currents were recorded using borosilicate glass electrodes polished to resistances of 14-23 MΩ using the cilium patch method described previously^50^. Single channel currents measured in the on-cilia configuration were recorded from PKD1^Null^:PKD2^Null^ HEK cells stably expressing the *in vivo* ARL13-GFP fluorescent cilia reporter that had been transiently transfected with PKD2 expression constructs. All our solutions were formulated with ultrapure water (Milli-Q IQ 7005 water). Pipette electrode solution contained (in mM): 140 NaCl, 10 HEPES, 1.8 CaCl_2_; pH was adjusted to 7.4 with NaOH. Bath solution contained (in mM): 120 KCl, 20 NaCl and 10 HEPES, 1.8 mM CaCl_2_; pH was adjusted to 7.4 using KOH. All solutions were osmotically balanced to 300 (±7) mOsm with d-mannitol. Data were collected using an Axopatch 200B patch-clamp amplifier, Digidata 1550B and pClamp 10 software. Single channel currents were digitized at 50 kHz and low pass filtered at 10 kHz. Data were analyzed by Igor Pro 7.00 (Wavemetrics).

### PKD2 cell-free synthesis, vesicle reconstitution and single channel recordings

Cell-free expression (CFE) methods for expressing PKD2 orthologues from yeast and human and reconstituting the channel proteins into lipid vesicles for functional characterization have been previously reported^30,51,52^. Lipids DPhPC (95%) and cholesterol (bovine, 5%) were obtained from Avanti Polar Lipids (Alabaster, USA) and mixed in chloroform, where vacuum evaporation on glass formed small unilaminar vesicles (SUVs), rehydrated and passed through a 100-nm polycarbonate filter with a Mini-Extruder (Avanti Polar Lipids). Cell-free protein of PKD2 was produced from a PKD2 pET19b plasmid under a T7 promoter with PURExpress In Vitro Protein Synthesis Kit from New England Biolabs, Inc (Ipswich, MA, USA). Diluted SUVs (1:500) were added to the reaction. GUVs were form from PKD2 incorporated SUVs on indium tin oxide-coated glass slides using a Vesicle Prep Pro (Nanion Technologies, Munich, Germany) under the standard electroformation program. GUVs reconstituted with PKD2 protein were used for electrophysiology experiments the same day. All GUV patch electrodes were made using borosilicate glass electrodes and were fire polished to resistances greater than 5–10 MΩ. Recording solutions for consisted of symmetrical recording solutions with 140 mM NaCl, 10 mM HEPES, and 300 mM glucose using a holding potential of −40 mV (Molecular Devices, San Jose, CA, USA, Axopatch 200B amplifier). Recordings were digitized with the Digidata 1550B (Molecular Devices) at 25 kHz and low pass filtered at 5 kHz. Recordings were analyzed with ClampFit v11.2.0.59 (Molecular Devices, San Jose, CA, USA) and IGOR Pro 9 (WaveMetrics, Portland, OR, USA). As a predetermined criterion, data was excluded from analysis when seal resistance fell below 5 MΩ due insufficient voltage control of the patched membrane.

### Isolation of murine embryonic fibroblasts and cell culture assays

To prepare mouse embryonic fibroblasts (MEFs), timed-pregnant female mice (E12.5) were euthanized, and the uterine horns containing embryos dissected. After releasing the embryos from the placental membranes, the heads and internal organs were removed, and the remaining embryonic tissue digested in using digestion buffer (containing 2mg/ml Collagenase Type 2 (17101015, Thermo Fisher Scientific) and DNase) in DMEM to generate a single cell suspension. The digestion was stopped by adding MEF culture medium, and the suspension seeded into 100-mm tissue culture plates and allowed to grow in growth medium (DMEM supplemented with 10% FBS (Gibco, cat. no. 10270-106), penicillin (100 U ml–1)/streptomycin (100 mg ml–1) (Gibco, cat. no. 15140-122) and 2 mM L-glutamine (Gibco, cat. no. 25030-024)) at 37 °C and in a humidified incubator5% (v/v) CO_2_. <u>Low temperature and protein stability assay</u>: Pkd2 D509V stability was assessed at low temperature. Briefly, MEF cells at passages P1-P2 were incubated for 48h in a humidified incubator (5% CO₂) at 27°C, 33°C or at 37°C. <u>Proteosome and</u> <u>lysosome inhibition assays</u>: MG132 (10 µM) (Selleckchem, Catalog # S2619) and Chloroquine (50 μM and 100 μM) (Sigma, Catalog # PHR1258) were added to MEFs growth medium for 6 and 24 hours respectively.

### Generation of the *Pkd2^D509V^* mouse strain

Signal guide RNAs (sgRNAs) for the *Pkd2* D509V missense mutation (5′-CAGGAGTTTCTGGAATTGTC -3) were designed using the CRISPR Design tool (http://crispr.mit.edu/). To edit the mouse genome, single-stranded oligodeoxynucleotide (ssODN) donor templates for the point mutation were synthesized (IDT). One-cell C57BL/6J embryos (Jackson Laboratories, # 000664) were generated using previously described protocols^53,54^. Embryo electroporation was performed using a protocol previously described with minimum changes^53^. Briefly, the electroporation solution was prepared stepwise, starting with equal volumes of Cas9 protein (1000 ng/µl stock, IDT, Catalog #10007806), tracrRNA (20 µM stock, IDT, Catalog #1072533), and sgRNA (20 µM stock, IDT), combined to make a 9 µl ribonucleoprotein (RNP) particle solution. The solution was prepared using RNase-free injection buffer (10 mM Tris-HCl, pH 7.4, 0.25 mM EDTA) and placed on ice for 10 minutes. The RNP + ssODN solution was prepared by adding 1 µl of ssODN (2000 ng/µl stock, IDT) to the RNP solution and allowing it to equilibrate at room temperature. The final electroporation mix was made by adding 35 µl of Opti-MEM (Gibco, 31985062) to the RNP + ssODN solution at room temperature. Embryos were electroporated by placing them in a 5 mm gap slide containing 45 µl of electroporation solution, in groups of up to 100 embryos. The following parameters were used for electroporation: pore pulse (voltage: 225 V; pulse length: 1.0 msec; pulse interval: 100 msec; number of pulses: 4; decay rate: 10%; polarity: +), transfer pulse (voltage: 20 V; pulse length: 50 msec; pulse interval: 50 msec; number of pulses: 5; decay rate: 40%; polarity: ±). Following electroporation, embryos were transferred into pseudopregnant CD1 females (Charles River, #022) using previously described protocols^53,54^. Founders were identified by PCR amplification of genomic DNA from tail biopsies, followed by sequencing of the PCR products^55^. The mutation was transmitted to the germline by breeding founders with wild type C57BL/6J females (Jackson Laboratories, # 000664).

### Genotyping analysis

Genomic DNA was isolated from mouse/embryos tails/toes and genotyped using the following primers allele-specific: For *Pkd2^D509V^* we used Pkd2-509V-F: 5’-CATAGTGCAGGGACTGACCA-3’ and Pkd2-509V-R 5’-TCTCCAACACTGAAATCGAAGA -3’ primers. The PCR reaction was carried out with an annealing temperature of 59°C for 35 seconds, over 35 amplification cycles. This PCR strategy amplifies a 412 base pair (bp) product from both the wild-type and mutant alleles. However, the D509V mutation introduces a restriction enzyme recognition site, which allows the mutant allele to be differentiated from the wild-type allele. Specifically, the mutation generates a new restriction enzyme site for BtsCI, resulting in the cleavage of the 412 bp product into two smaller fragments: 300 bp and 112 bp. After amplification, the PCR products were digested with BtsCI (20 U per reaction) (New England Biolabs, Catalog # R0647S) at 50°C for 60 minutes. Following digestion, the reaction was heat-inactivated at 80°C for 20 minutes to stop the enzyme activity. This PCR generates a 412bps band in wild types (Pkd2^+/+^), three bands of 412bps+300bps+112bps in heterozygous *Pkd2^+/D509V^* mice and two bands of 300bps+112bps in homozygous mutants *Pkd2 ^D509V/D509V^*. For *Pkd2^fl/fl^; Pax8^rtTA^; Tet-O-Cre* genotyping was performed using the following primers: Pkd2flox11-13(A)-F 5’-CCTTTCCTCTGTGTTCTGGGGAG-3’, and Pkd2flox11-13(B)-R 5’-GTTTGATGCTTAGCAGATGATGGC-3’ to identify the *Pkd2^f/f^*allele; oIMR7385 5’-CCATGTCTAGACTGGACAAGA-3’ and oIMR7386 5’-CTCCAGGCCACATATGATTAG-3’ to identify the presence of the Pax8^rtTA^ transgene and Cre-200F: 5’-AGGTTCGTTCACTCATGGA-3’ and Cre-200R: 5’-TCGACCAG TTTAGTTACCC-3’ to identify Cre positive and negative mice. PCRs were performed under the following conditions: an annealing temperature of 56°C for the *Pkd2^f/f^* allele, 63°C for the *Pax8^rtTA^* transgene, and 59°C for the *Tet-O-Cre* transgene, with 35 cycles for each. The PCR products were resolved on 2% NuSieve agarose gels. The expected product sizes were approximately 318 bp for the Pkd2^fl^ allele, 595 bp for the *Pax8^rtTA^* allele, and 235 bp for the Cre allele.

### *Pkd2* inactivation

The *Pkd2* conditional allele (*Pkd2^fl/fl^*) used in these studies has been previously described^13^. The *Pkd2^fl/fl^; Pax8^rtTA^; Tet-O-Cre* mouse model is an orthologous mouse model of ADPKD that uses the murine *Pax8* promoter to direct expression of the reverse tetracycline-dependent transactivator (rtTA) to all renal tubular compartments (proximal and distal tubules as well as collecting ducts)^56^. We crossed *Pkd2^fl/fl^; Pax8^rtTA^; Tet-O-Cre* males with *Pkd2^+/D509V^* heterozygous females to generate *Pkd2^D509V/fl^; Pax8^rtTA^; Tet-O-Cre* positive mutant and *Pkd2^D5^*^09^*^/fl^; Pax8^rtTA^; Tet-O-Cre* positive controls. To induce *Pkd2* deletion, experimental mice were treated with a single daily intraperitoneal (IP) injection of Doxycycline Hyclate (Sigma-Aldrich, D9891) 50 mg/Kg body weight, diluted in sterile distilled water at P10 and P11. Littermate controls were treated with Doxycycline in the same manner as experimental mice. All mice were maintained on an inbred C57BL/6 background. Mice of both sexes were included in this study.

### Histology, immunocytochemistry, and confocal microscopy

The kidneys from transgenic mice were fixed in 4% paraformaldehyde for 24 h, embedded in paraffin sectioned on a microtome (5µm thickness), mounted on glass slides, and stained with hematoxylin and eosin (H&E) or Dolichos Biflorus Agglutinin (DBA) and fluorescein-labeled Lotus Tetragonolobus Lectin (LTL). <u>Cystic Index calculation</u>: Images were photographed by a digital camera (Infinity 2) connected to a Nikon eclipse E600 upright microscope. The total kidney area and cystic and non-cystic areas were measured using ImageJ (Fiji, Madison, WI). The cystic index is expressed as the percentage of the cystic area or lumen versus the total area of the kidney. <u>Cyst Origin Assessment</u>: was performed by staining paraffin kidney sections permeabilized with 0.1% Triton X-100, and blocked with 1% bovine serum albumin and 5% fetal serum bovine in PBS using rhodamine-labeled Dolichos Biflorus Agglutinin (DBA) and fluorescein-labeled Lotus Tetragonolobus Lectin (LTL) (Vector Laboratories, #RL-1032 and #FL-1321 respectively). Nuclei were counterstained with DAPI, washed three times with PBST, and mounted with Fluoromount-G™ Mounting Medium (Invitrogen, #00-4958-02). Images were photographed using a Nikon W-1 spinning disk confocal microscope at the UMB-SOM Confocal Microscopy Core-Baltimore, Maryland.Immunocytochemistry of sectioned kidneys used for the visualization of collecting duct primary cilia were fixed with 4% paraformaldehyde, permeabilized with 0.2% Triton X-100, and blocked by 10% bovine serum albumin in PBS. Primary antibodies used were 1:1000 anti-ARL13B (Protientech, 17711-1-AP) and 1:500 anti-aquaporin 2 (Abcam, EPR21080). Cell nuclear visualized with DAPI nuclear staining (Invitrogen) in some experiments. Cultured PKD1^null^:PKD2^null^ HEK cells expressing human ARL13B-6Gly-GFP variant channels were fixed with 4% paraformaldehyde (PFA), permeabilized with 0.2% Triton X-100, and blocked by 10% bovine serum albumin in PBS. Cultured HEK cells and tissue were mounted on glass slides and treated with Fluoshield from Sigma-Aldrich (St. Louis MO). Primary cilia localization of the HA-PKD2 variant channels expressed in HEK PKD1^Null^:PKD2^Null^ cells were visualized by [1/1500] by treatment with 594-conjugated HA tag polyclonal antibodies (protientech, CL594-51064). Confocal images were obtained using an inverted Nikon A1 with a 60x silicon oil immersion, 1.3 N.A. objective. Confocal images were further processed with FIJI ImageJ (National Institutes of Health). Cilia lengths measured by masking the ARL13B signal with minimum 1μm threshold of cilia length. Super-resolution images using the SIM method were captured under 100× magnification using the Nikon Structured Illumination Super-Resolution Microscope with piezo stepping. 3D-SIM images were further processed with Imaris software (Oxford Instruments). All histologic analyses were performed by blinded observers.

### Western Blot analysis

Frozen specimens (embryos) and MEF cells were lysed using Cell Lysis Buffer [20 mM Na-P (pH 7.2), 150 mM NaCl, 1mM EDTA, 10% Glycerol, 1% Triton-X100] and a cocktail of protease inhibitors (Sigma, #P8340, 1:100). Tissues and cell lysates were homogenized with a motor driven Polytron or pipetting up and down respectively, incubated for 40 minutes at 4°C rotating in lysis buffer and centrifuged at 14k rpms for 30 minutes at 4°C. The supernatant was transferred to a fresh tube and the protein concentration was measured using the BCA protein assay kit (Pierce). Fifty micrograms of protein were loaded on NuPAGE™ 3-8%, Tris-Acetate mini protein gels (Invitrogen™, #EA03755BOX). Proteins were blotted using an anti-rabbit PKD2 antibody (provided by the PKD-RRC), vinculin antibody (Sigma, #V9131, clone hVIN-1; 1:2000) and β-Actin antibody (Cell Signaling, clone 13E5-Rabbit #4970, 1:1000) and secondary HRP-linked whole anti-mouse and anti-rabbit antibodies, (Amersham, NA-934, 1:10000). Blots were imaged using a GelDoc Go imaging system (Bio-Rad) and band density was quantified using ImageJ (Fiji, Madison, WI).

### Transcriptomic RNA sequencing

Embryonic kidneys were isolated at E16.5 from wild-type control mice and homozygous mice carrying the Pkd2-D509V mutation. Pregnant dams were euthanized according to approved animal-use protocols, embryos were rapidly harvested, and kidneys were microdissected in cold sterile PBS to minimize RNA degradation. Dissected kidneys were immediately transferred into RNase-free tubes, snap-frozen or stabilized in RNA-preservation reagent and shipped on dry ice to Plasmidsaurus (South San Francisco, CA) for RNA processing and transcriptomic analysis. Transcriptomic profiling was performed using the 10x Genomics platform, with Plasmidsaurus generating sequencing libraries and performing transcriptomic read acquisition. Resulting sequencing data from WT and homozygous D509V kidneys were used for downstream differential gene-expression analysis to identify transcriptional programs altered by the D509V variant, with particular attention to pathways related to PKD2 function in primary cilia biology, epithelial differentiation, kidney development, and cystic disease progression.

### Statistical Analysis

Statistical methods used to determine significance are described in corresponding figure legends and reported as *P* values. Study design sample sizes were determined by variant analysis of previouly published data sets. Briefly, electrophysiology and imaging datasets were analyzed (GraphPad or Origen) using one way ANOVA, Student’s t-tests two tailed paired (equal sample sizes) or unpaired (unequal sample sizes).

### Animal Welfare Statement

Animals were housed at AALAS certified facilities located at University of Maryland, School of Medicine, Feinberg Medical School, Northwestern University and Johns Hopkins University. All animal procedures and protocols were approved by the Institutional Animal Care and Use Committees (IACUCs).

### Materials availability statement

All mammalian cell expression constructs used in this study are available without restriction upon written request to the corresponding author. Transgenic animals are available from the Polycystic Kidney Disease Research Resource Consortium (PKD RRC).

## RESULTS

### PKD2 D511V variant impact on channel stability and structure

We began our study by examining the biochemical and structural biology consequences of the D511V variant on PKD2 channel stability and structure. We used the BacMan system to transiently express WT and D511V channels (residues 53–792) with an N-terminal maltose binding protein (MBP) fusion tags in HEK 293S GnTI− suspension culture cells. Channel protein was affinity purified with amylose resin and the MBP tag was removed by TEV protease cleavage and separated by size exclusion chromatography (SEC, Superose 6 column). The elution peak fractions corresponding to PKD2 homotetramer were pooled and concentrated for thermostability analysis and subsequent cryo-EM structural determination. Using fluorescence detection size exclusion chromatography (FSEC) to test channel stability by tryptophan fluorescence over temperature gradients, the WT PKD2 showed a clear thermal transition whereas the D511V variant exhibits multiple denaturing intermediates, indicating a global destabilization of the channel tetramer was caused by the VSD mutation (**Figure 1 A-B**). After plunge freezing the D511V protein reconstituted in detergent micelles and collecting cryo-EM data sets using state of the art electron microscope, we observed both assembled and disassembled channels in micrographs. More than two million single particles of the variant channels were collected, processed, and refined to resolve the homomeric PKD2 D511V structure at 3.1 Å overall resolution (**Figure 1C**, **Supplemental Figure 2, Supplemental Figure 3**). The VSD of the D511V channel structure was captured in its deactivated state (down state), and with the internal gate of the ion-conducting pore domain closed (**Figure 1D**). Compared to our previously determined WT structure, the global domain-swapped assembly and overall domain architecture of the variant channel was retained. However, the D511 residue normally found in the S3 within the WT VSD was replaced by a valine due to the missense substitution. Normally, the D511 side chain forms anions interactions with the carboxylate side chains of positively charged lysine residues (K572, K575) of the S4 (**Figure 1E, Supplemental Figure 1**) ^8,29^. These so called *‘gating charge’* salt-bridges are among the most highly conserved interactions within voltage-gated channels but are exclusive to polycystin subfamily members within the transient receptor potential (TRP) channel family (**Supplemental Figure 1B**)^57–59^. The D511 residue is present in all 3 members of polycystin subfamily and is conserved in homomeric prokaryotic voltage-gated sodium (Na_v_) and eukaryotic potassium channels (Kv) (**Figure 1E, Supplemental Figure 1A**). Since eukaryotic Na_v_s and Ca_v_s are single pseudotetrameric polypeptides that likely evolved from an ancestral prokaryotic homomeric channel after several rounds of gene duplication, the gating charge-anion charge interactions within their VSDs are also conserved across phyla^60–62^. Evidently, disrupting this highly conserved transmembrane interaction with a neutralizing missense mutation is sufficient to drastically destabilize the nascent PKD2 channel protein. Accordingly, we observed more broken and aggregated particles in D511V cryo-EM 3D classifications (class 2 and 3 in **Supplemental Figure 4**) than in WT preparations. Contrast to the WT PKD2 or TOP domain variants (C331S, R322Q/W R325Q/P) structures that we solved previously^25,47^, the initial particle alignment converges from a well-ordered TOP domain, which enables structural model all TOP domain fingers and crucial N-glycans beyond monosaccharides. Not surprisingly, the lower half of the VSD is the least resolved, suggesting VSD mutation D511V disrupted the stability of this region (**Supplemental Figure 2E**). Next, we will evaluate the consequences of this variant on channel trafficking and function within the primary cilia membrane.

**Figure 1.**
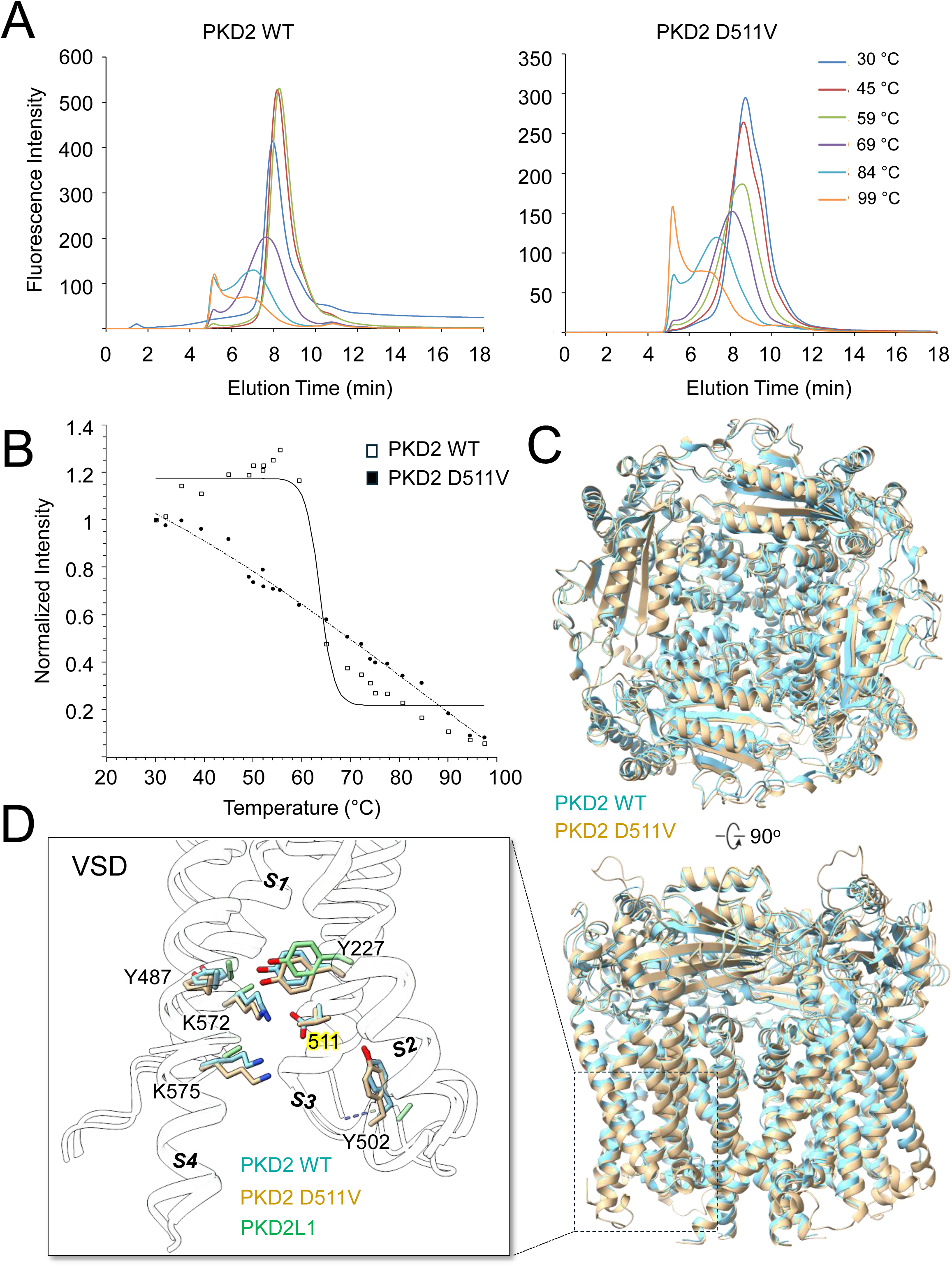
D511V variant destabilizes PKD2 by neutralizing gating charge interactions within the VSD. **A)** Size-exclusion chromatography (SEC) profiles of the WT and D511V channels after heating. **B)** Fluorescence size-exclusion chromatography (FSEC) thermal denature profiles for the PKD2 channels. While a thermal transition temperature at 64.0 +/- 0.9 °C was observed for wild type PKD2, no clear single transition temperature was observed under current experimental condition. **C**) Cryo-EM structure of the PKD2 D511V variant channel solved at 3.1 Å. *Top*, external and transmembrane view and structural alignment of the WT and D511V channels **D**) Structural alignment of the voltage sensor domain found in polycystin channels (PKD2 WT PDB 6T9N, D511V, PKD2L1 PDB 6DU8)^32,90^.

### PKD2 D511V variant abolishes channel primary cilia trafficking and function in membranes

Although the D511V variant destabilized the PKD2 channel structure, a small percentage still assembled into tetrameric particles— suggesting that they retain some capacity to traffic and function in cells. While previous work reconstituting ER PKD2 protein into lipid bilayers suggests that the D511V mutation causes a loss of function, it is unknown how this variant affects the biosynthesis of the mature population of these channels in primary cilia membranes^63^. Using structured illumination microscopy (SIM), we first compared primary cilia trafficking of exogenously expressed channels using immunofluorescence in PKD1 and PKD2 null HEK cell line (PKD1^null^:PKD2^null^), which rules-out the contribution of endogenous polycystins^48,49^. We then stably expressed ARL13B-GFP to visualize the primary cilia compartment and N-terminal HA-tagged channels labeled with anti-HA antibodies (**Figure 2A**). Imaging analysis of fixed cells demonstrated abundant WT channel trafficking to the primary cilia, as assessed by Pearson’s coefficient analysis with the ciliary ARL13B signal (**Figure 2B**). In contrast, the D511V variant channel exhibited defective ciliary trafficking with significantly shorter primary cilia compared to those expressing WT (**Figure 2A, C**). To confirm the absence of ciliary trafficking by the D511V variant and to evaluate its potential functional impact caused by the VSD mutation, we conducted cilia electrophysiology (**Figure 3A, B**)^23,24^. After establishing high-resistance seals from cells expressing the WT PKD2, single channel open events were present in 83% of cilia patches (15/18), with three openings per patch being the most frequent (6/18) channel density observed in the experiment (**Figure 3C**). In contrast, despite forming high resistance seals and voltage-clamping more than 23 cilia membranes, no single channel events were detected from cells expressing the D511V variant (0/23). These electrophysiology results support the previous biochemical and imaging results, suggesting the D511V mutation disrupts VSD salt bridges destabilizing channel assembly and abolishes cilia trafficking. To test this hypothesis further, we introduced a glutamate substitution at the same site (D511E) which preserves the proposed side chain counter charge interaction with the K575 gating charge. As expected, the D511E mutation traffics to the primary cilia and retained channel functional activity— abet with a reduced numbers (**Figure 2A, C**, **Figure 3A-C**). To further test if the D511V variant can produce active channels, we used a combined cell-free expression and giant unilamellar vesicle reconstitution approach (CFE+GUV), a method that has been successfully applied to functional assays of yeast and human PKD2 orthologues (**Figure 3D**)^30,51,52^. Activity from CFE-derived PKD2-GFP protein reconstituted in GUVs comprised of DPhPC was enhanced with the addition of 5% cholesterol, as determined by the increased maximum number of open channel events and reduced number of empty patches (**Figure 3E, F**). However, under the same conditions, no D511V channel events were detected in the voltage clamped GUVs membranes. Together, these in vitro findings show that D511V profoundly destabilizes PKD2, yielding channels that fail to traffic to primary cilia and remain completely non-functional even when forcibly reconstituted into lipid membranes. We therefore hypothesized that this combined loss of protein stability, ciliary trafficking, and channel activity would produce significant ADPKD pathology in vivo, which we assess in the next section.

**Figure 2.**
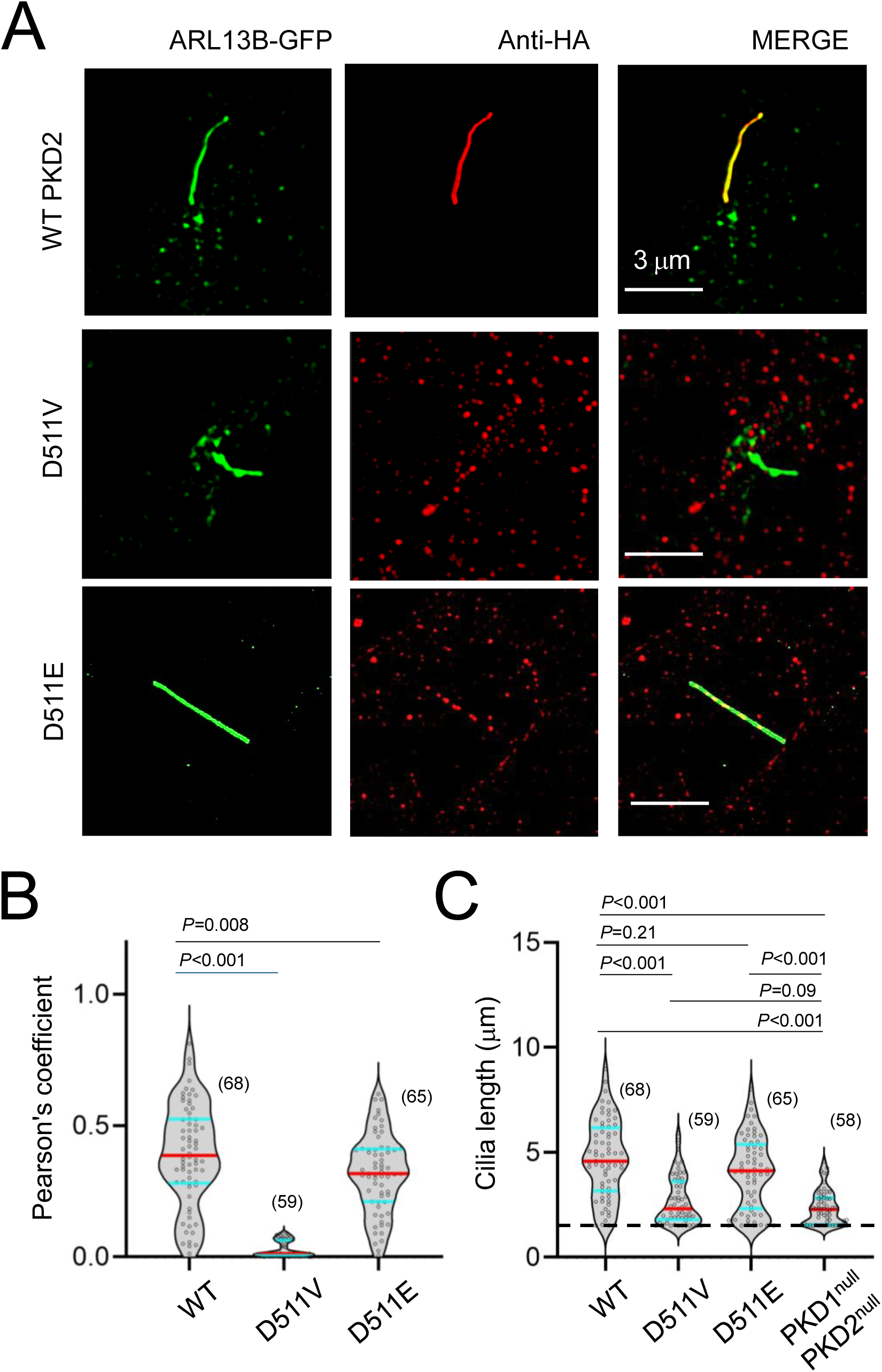
D511V variant abolishes PKD2 primary cilia trafficking. **A**) Super resolution SIM (structured illumination microscopy, Scale bar = 3 μm) images of HEK PKD1^Null^:PKD2^Null^ cells stably expressing ARL13-GFP to visualize the primary cilia. HA-PKD2 WT and variant channels immunolabeled with anti-HA. **B**) Violin plots of cilia-channel fluorescence colocalization analysis using the Pearson’s correlation coefficient. Red lines indicate the median and blue lines are equal to quartiles. **C**) Comparison of cilia length from the indicated transfected HEK cell lines. *P*-values indicate Student’s t-test results (see methods) comparing WT and variants channels (*P* < 0.05). Sample size (cilia) for each group are indicated in the parenthesis and error bars are equal to standard deviation.

**Figure 3.**
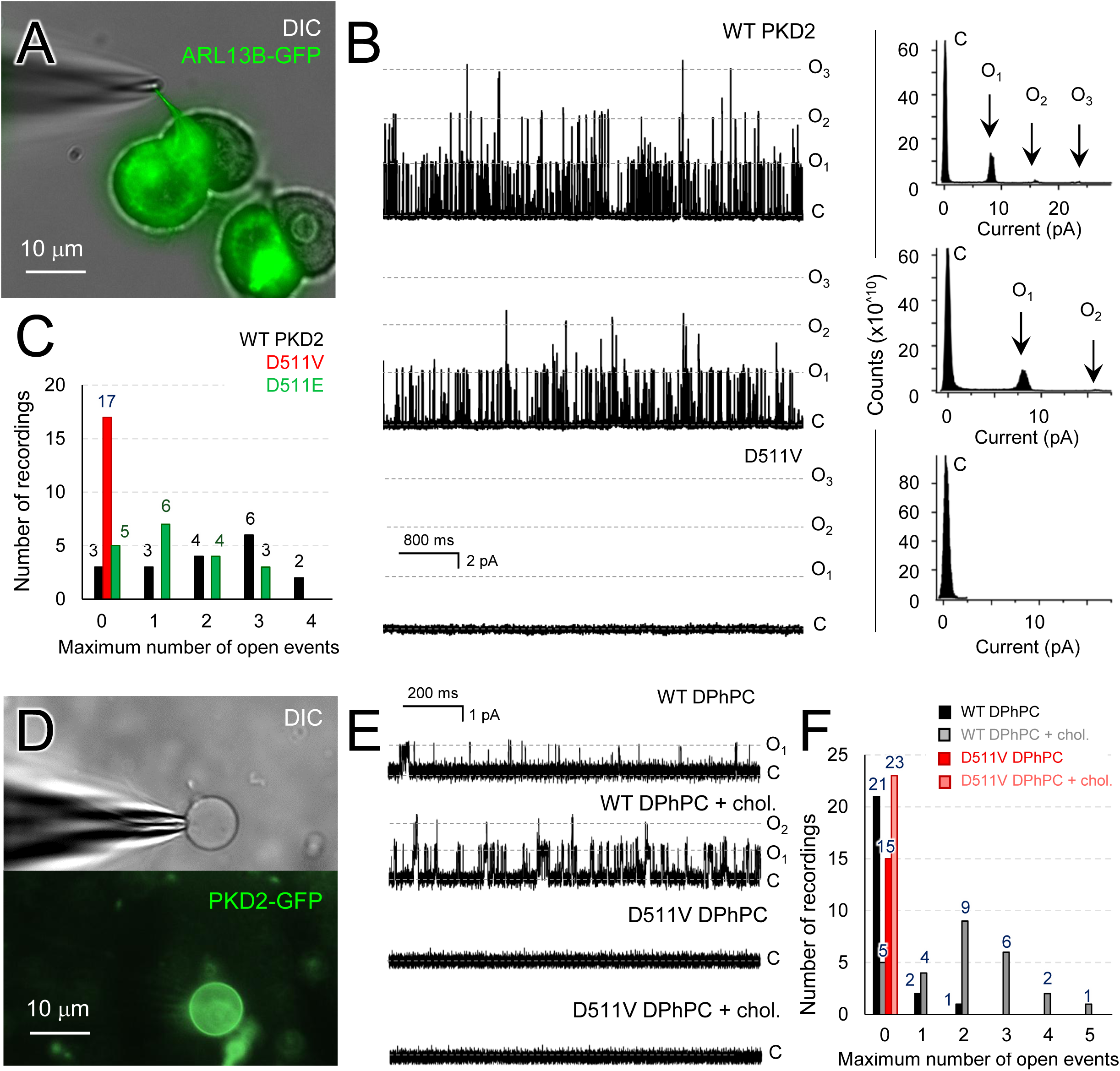
PKD2 D511V variant is nonfunctional following vesical reconstitution and is absent from the activatable channel population in primary cilia membranes. **A**) *Left*, image of a voltage clamped primary cilia in the on-cilia configuration. The primary cilium of HEK (PKD1^Null^:PKD2^Null^) cells transiently expressing PKD2 channels are illuminated by ARL13B-GFP under 475 nm laser excitation. *Right*, example single channel recordings from the cilia membrane activated by depolarizing voltage steps. Note, no open channel events were observed in cilia patch recordings from cells expressing D511V channels. **B**) *Left*, example polycystin unitary single channel events recorded from cilia membranes. *Right*, corresponding histograms indicating the frequency of a variable simultaneous open channel events (O_1_-O_3_) and closure (C). Note, no channel openings are detected from cilia recordings from cells expressing D511V channels. **C**) Column plots indicating the frequency of the maximum simultaneous open channel events from cilia patches. Number of events are indicated in parenthesis. **D**) Image of a voltage clamped giant unilaminear vesicals (GUVs) reconstituted with cell free expressed PKD2-GFP protein. **E**) *Top*, example WT PKD2-GFP unitary single channel events recorded (HP = 40mV) from GUVs composed of either 1,2-diphytanoyl-sn-glycero-phosphocholine (DPhPC) or 95% DPhPC and 5% cholesterol (Bovine). *Bottom*, lack of single channel events from D511V channels reconstituted in GUVs. **F**) Column plots indicating the frequency of the maximum simultaneous open channel events from GUVs patches. Number of recordings indicated above for each column.

### PKD2 voltage sensor variant expression causes in vivo lethality and primary cilia destabilization

Our *in vitro* data provide compelling evidence that the ADPKD-associated D511V variant results in complete channel dysfunction that arises from the destabilization of VSD salt-bridge interactions that impairs channel assembly and prevents trafficking to the primary cilia of kidney cells. To explore the cystogenic potential of this variant within a murine context, we used CRISPR/Cas9 gene-editing technology to generate a mouse model (*Pkd2^D509V^*) that carries a germline mutation equivalent to the human D511V variant (**Figure 4A, Supplemental Figure 1**). Heterozygous (*Pkd2^+/D509V^*) mice are viable, healthy and fertile, with no apparent phenotypes at approximately 6 months of age. To determine the viability of homozygous mutants, we intercrossed *Pkd2^+/D509V^* males and females. Genotypic analysis of their offspring at postnatal day P0 confirmed the expected presence of wild type *Pkd2^+/+^* and heterozygous *Pkd2^+/D509V^* progeny, however, no viable homozygous *Pkd2^D509V/D509V^* pups were detected at this stage (**Supplemental Table 2**). To investigate whether the absence of viable *Pkd2^D509V/D509V^*offspring was due to embryonic lethality, we conducted timed-pregnancies studies using *Pkd2^+/D509V^* heterozygous breeders. Pregnant females were sacrificed, and embryos collected at embryonic day E14.5. Genotypic analysis at this stage identified the presence of homozygous embryos, albeit at slightly reduced Mendelian ratios (**Supplemental Table 2**). Viable *Pkd2^D509V/D509V^* embryos exhibited several phenotypic abnormalities including polyhydramnios, edema and occasional hemorrhage (**Figure 4A-B**) —suggesting that the mid-gestation lethality was due to cardiovascular defects— features that are consistent with the phenotype observed in *Pkd2* null embryos and other *Pkd2* strains with mutations preventing the proper trafficking of Pkd2 protein to the cilia^10,17,18^.

**Figure 4.**
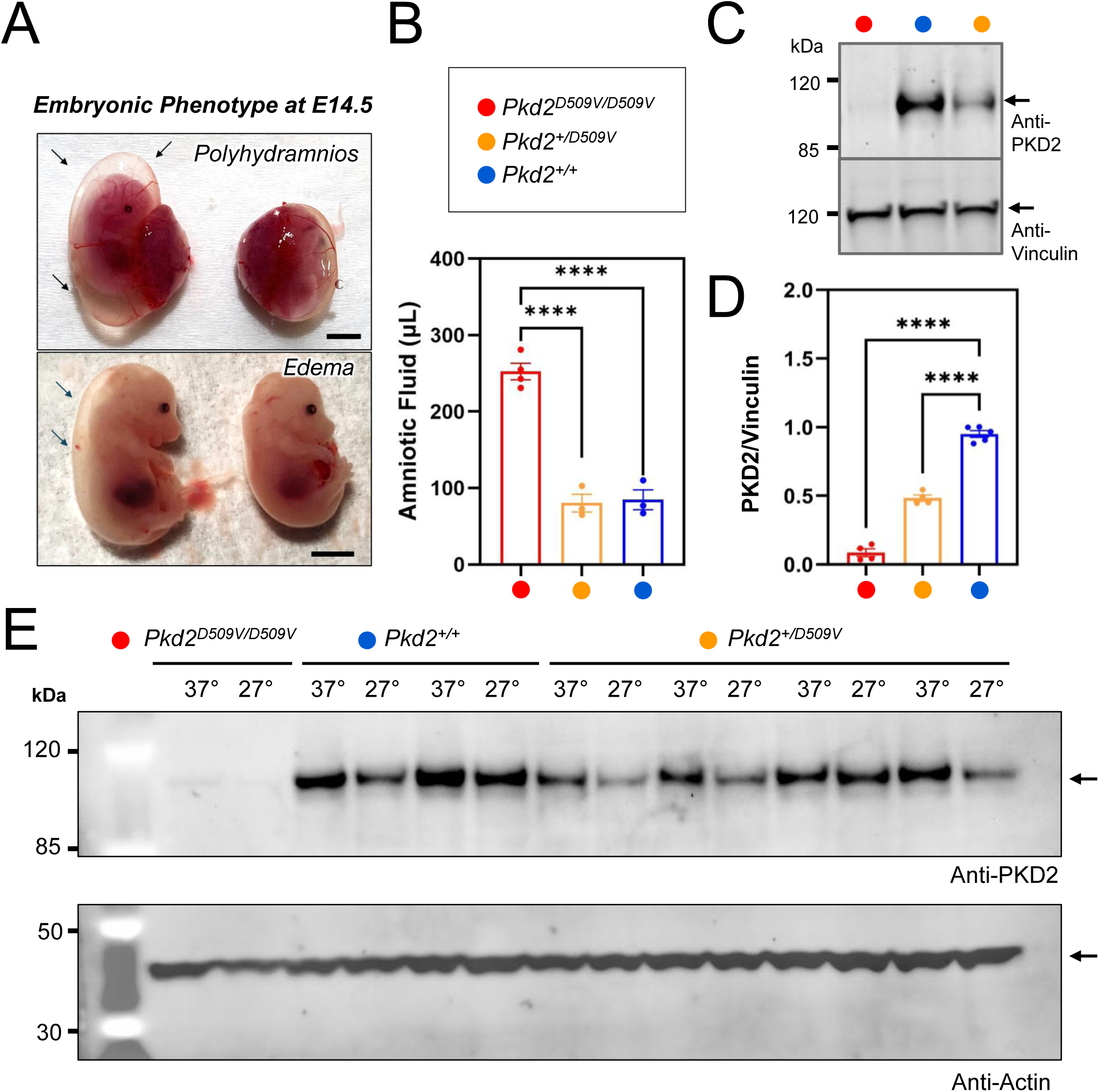
Mice expressing homozygous *Pkd2^D509V^* variant alleles are embryonic lethal. **A**) Gross appearance of homozygous *Pkd2^D509V/D509V^*(red circle) and *Pkd2^+/+^* control (blue circle) littermate embryos harvested at E14.5. Homozygous embryos exhibit severe polyhydramnios (black arrows, upper panel). In the lower panel, after removing the yolk sac, the homozygous embryos show the presence of edema (black arrows). We observed that the majority of embryos die at mid-gestation (∼E14.5–E15.5) and display a phenotype similar to that seen in *Pkd2*^-/-^ null embryos (data not shown), suggesting that the embryos may die due to cardiovascular complications, as observed in null embryos. The scale bar represents 2 mm. **B**) Quantification of amniotic fluid demonstrated that homozygous mutants develop polyhydramnios. Data are presented as mean ± SEM, with comparisons performed using one-way ANOVA followed by Tukey’s multiple comparisons test in GraphPad Prism 9.0 software. **C**) Western blot showing expression of polycystin protein from *Pkd2^D509V/D509V^* embryos (E14.5), *Pkd2^+/D509V^* and *Pkd2^+/+^* littermates. Note the dramatic reduction of protein expression in homozygous embryos. Vinculin was used as loading control. **D**) Quantification of polycystin protein expression normalized to vinculin in homozygous, heterozygous and wild-type embryos. Data are presented as mean ± SEM, with statistical comparisons performed using one-way ANOVA followed by Tukey’s multiple comparisons test. **E**) Western blot analysis of Pkd2 protein expression from MEF cells isolated from homozygous (red), wild-type (blue) and heterozygous (orange) embryos cultured at 37°C and 27°C (to rescue thermodynamic instability) in a 5% CO₂ incubator for 48 hours. Actin was used as a loading control. Notably, expression levels of both wild-type and variant *Pkd2^D509V^* remain unchanged after culturing cells at 27°C, suggesting that the variant is not a temperature-sensitive folding mutant.

To determine whether variant Pkd2 variant alleles affected protein expression, we performed western blot analysis, using cell lysates derived from homozygous *Pkd2^D509V/D509V^* and control littermates (*Pkd2^+/D509V^* and *Pkd2^+/+^*) embryos at E14.5, and probed with an antibody against Pkd2 protein and vinculin (as loading control) (**Figure 4C**). We observed a dramatic reduction of Pkd2 protein expression in homozygous *Pkd2^D509V/D509V^* compared to littermate controls *Pkd2^+/+^*. Interestingly, heterozygous *Pkd2^+/D509V^* embryos exhibited also approximately a ∼50% reduction of Pkd2 expression, consistent with the presence of only one functional allele (**Figure 4C-D**). Based on the results from overexpressed VSD variant in cell lines, we sought to the D509V mutations allelic impact on renal primary cilia morphology. Embryonic kidneys from littermates expressing variant alleles were harvested, fixed, section and immunolabeled with anti-ARL13B antibodies to visualize the primary cilia (**Supplemental Figure 5A**). Primary cilia from mice expressing D509V alleles were significantly shorter, but the greatest impact was observed in the tissue from homozygous mice (**Supplemental Figure 5B**). These findings suggest that the missense variant D509V leads to a significant loss of PKD2 expression and cilia stability *in vivo*, contributing to the severe phenotypes observed during embryonic development. To further understand cilia destabilizing effect, we carried out RNA sequencing of embryonic kidneys isolated from WT and *Pkd2^D509V/D509V^* mice (**Supplemental Figure 5C**). Transcriptomic analysis suggests a coherent down regulation of pathways that regulate microtubule, Ca^2+^ signaling, G-protein–dependent regulatory pathways required for maintenance of the primary cilium (Cdc42, Gnai2, Gnb1, Gnb2, Calm3, Dynll2, Tubb4b, Tubb2a, and Tuba1b). Conversely, increased Rhoa and Cltc expression may reflect compensatory cytoskeletal remodeling and altered clathrin-dependent membrane protein trafficking. Together these exploratory results support a model in which PKD2 channel dysfunction destabilizes the ciliary axoneme and ciliary membrane organization, contributing to loss of primary cilia in mutant kidney tissue.

Previous studies expressing the D511V variant in *HeLa* cells and in fruit flies (D. melanogaster) identified defective protein processing associated with this mutation ^64,65^. To further investigate this phenomenon, we conducted western blot analysis of murine embryonic fibroblasts (MEFs) isolated from *Pkd2^D509V/D509V^*mutant embryos and control littermates (*Pkd2^+/D509V^* and *Pkd2^+/+^*). We cultured MEFs cells at physiological (37°C) and lower temperatures (27°C and 33°C) to assess whether reduced heat could improve protein stability and expression (**Figure 4E, Supplemental Figure 6**), as characterized in our FSEC thermal denaturing results (**Figure 1A, B**). However, our analysis revealed a significant reduction of D509V protein, regardless of the culture temperature. Recognizing that proteins with aberrant folding are often targeted for degradation, we attempted a second approach to rescue the variant expression through inhibition of the lysosomal and proteasomal degradation pathways by treating MEF cultures with chloroquine (CQ) and MG132, respectively (**Supplemental Figure 7A, B**). However, neither reagent altered PKD2 variant protein levels, indicating that pharmacological inhibition of protein degradation pathways is insufficient to stabilize the variant protein and restore its expression. In summary, our findings demonstrate that the equivalent D511V mutations leads to a near-complete loss of PKD2 protein expression *in vivo*. This severe reduction in PKD2 protein status underscores the critical role of channel protein levels in maintaining cellular function and murine vitality.

### ADPKD mouse model based on the clinically-relevant PKD2 voltage sensor variant

The renal cystic phenotype observed in *Pkd2* null embryos at E14.5, the stage when fetal demise typically occurs, is generally mild or absent^18,66^. As a result, it is not feasible to evaluate the specific contribution of the D511V variant to cystogenesis under these conditions^10^. Thus, to further evaluate the cystogenic propensity of variant, we crossed *Pkd2^+/D509V^* mice with mice expressing the kidney-specific doxycycline-dependent *Pkd2* repression strain abbreviated *Pkd2^fl/fl^* (for *Pkd2^fl/fl^*; *Pax8^rtTA^; TetO-cre)*^19^. The resulting compound heterozygous mice, *Pkd2^D509V/fl^*, expresses the human variant equivalent mutation on one allele, while the otherwise functional *Pkd2^fl^* allele is repressible after doxycycline treatment. As expected, *Pkd2^D509V/fl^* mice were viable after birth. Two cohorts (*Pkd2^+/fl^* and *Pkd2^D509V/fl^)* of 10-11 day old mice were intraperitoneal injected (IP) with doxycycline (50mg/Kg) to inactivate the *Pkd2^fl^* allele. Mice were evaluated after 15 days (P25) using histology, kidney-to-body weight (KBW) and cystic index (CI) analysis (**Figure 5A**). In response to *Pkd2^fl^* inactivation, mice expressing the D509V mutation exhibited rapid and severe kidney enlargement and cystogenesis, whereas *Pkd2^+/fl^* mice showed no cyst development (**Figure 5B**). Quantitative assessments revealed kidney-to-body weight ratios (KBW) in doxycycline treated *Pkd2^D509V/fl^* mice was approximately six to ten times higher than in their control littermates, reflecting a significant renal enlargement, and the cystic index 3-4 times higher than in controls (**Figure 5C-E**), emphasizing the extensive cystogenesis driven by the D509V mutation.

**Figure 5.**
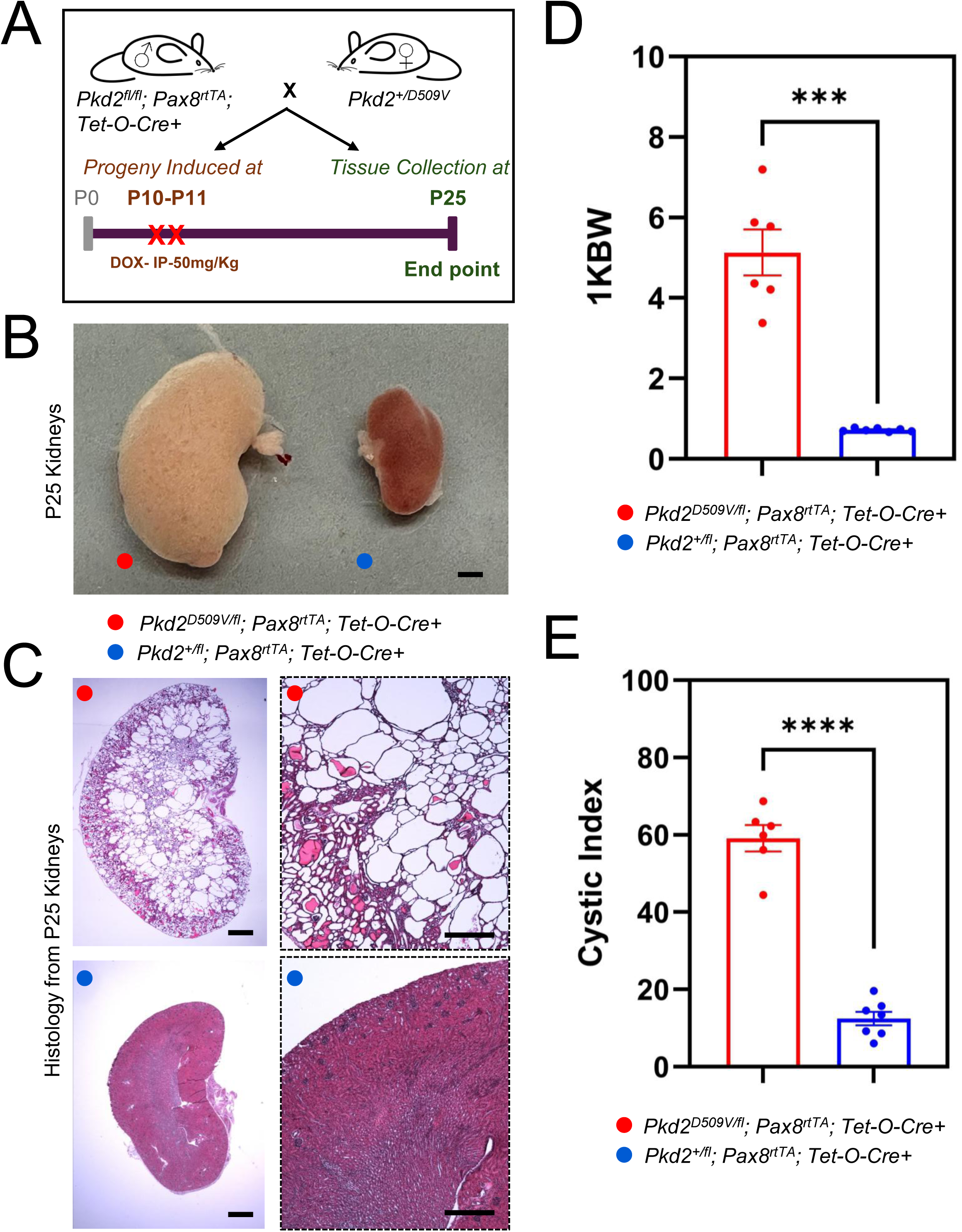
Compound transgenic mice expressing the analogous VSD PKD2 variant have rapid renal cyst development. **A**) Schematic showing the *in vivo* study design. We induced kidney-specific deletion of the *Pkd2* floxed alleles (*Pkd2^fl^; Pax8rtTA; Tet-O-Cre*) in compound transgetic mice expressing wild-type *Pkd2* (*Pkd2^+/fl^*) or the analogous D511V PKD2 VSD variant (*Pkd2^D509V/fl^*) at P10-P11 using doxycycline hyclate (50mg/Kg) via intraperitoneal (IP). Kidneys were harvested 15 days after *Pkd2^fl^* inactivation to evaluate cysts formation and disease progression. **B**) Gross appearance of kidneys (P25) from littermates expressing the variant *Pkd2^D509V^* allele (red circle) or wild-type *Pkd2^+^* allele. The scale bar = 1mm. **C**) H&E staining demonstring cortico-medullary cysts in kidneys harvested from *Pkd2^D509V/fl^*but not in kidneys from *Pkd2^+/fl^*. The scale bars represent 1mm and 500μm (left panel and right panel respectively). **D**) Kidney weight-to-body-weight ratio (1 kidney) 1KBW and (**E**) cystic index at P25. Data are presented as mean ± SEM and comparisons were performed using unpaired Welch’s corrected t-test with P-values <0.05 considered significant.

To determine the cellular origin of the renal cysts, we stained kidney sections from pups *Pkd2^D509V/fl^*and *Pkd2^+/fl^* at P25 after doxycycline induction (P10-P11) using specific nephron segment markers *Lotus* tetragolonobus lectin (LTL) for proximal tubules; and Dolichos biflorus agglutinin (DBA) for distal tubules. (**Figure 6A**). We observed cysts arising from both proximal and distal tubules, and occasionally, from other segments of the nephron in *Pkd2^D509V/fl^* kidney sections, but no cysts were observed in *Pkd2^+/fl^* control littermates, indicating a clear widespread impact of the D509V missense variant on the nephron structure. Since constitutive homozygous expression of the variant resulted in cilia degeneration during renal embryonic development (**Supplemental Figure 5B**), we examined its impact on renal primary cilia during disease progression. Two weeks after doxycycline treatment, primary cilia from renal tubular epithelium of *Pkd2^D509V/fl^* mice were less dense and significantly shorter compared to the untreated control group (**Figure 6B**). These findings suggest the VSD variant cystogenic mechanism is driven by a destabilizing structural defect, that causes hypomorphic PKD2 protein levels and promotes primary cilia degeneration across renal tubule segments. Furthermore, the established ADPKD animal model provides a valuable tool to investigate the downstream effects of PKD2 channel misfolding and attenuated gene-dosage. This experimental platform can be leveraged to evaluate potential therapeutic interventions aimed at restoring or correcting PKD2 which advances efforts toward effective treatment for polycystic kidney disease.

**Figure 6.**
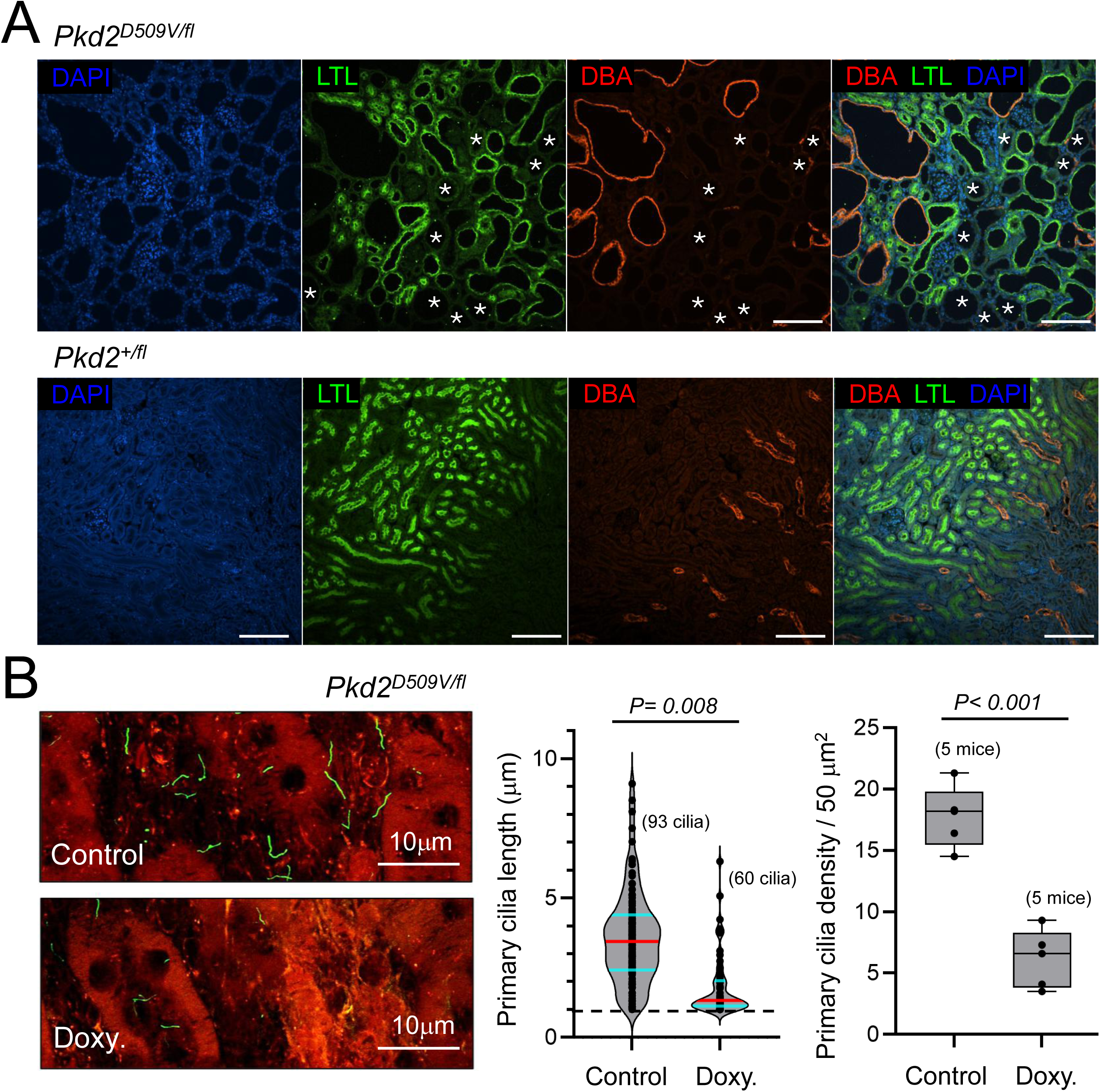
Kidney cyst origen and primary cilia destabilization in *Pkd2^D509V/fl^* compound heterozygous mice. **A**) Immunostained kidney sections from *Pkd2^D509V/fl^*and *Pkd2^+/fl^* mice after doxycycline induction as shown in Figure 4A. The distal tubules were labeled with *Dolichos biflorus agglutinin* (DBA) (red), while the proximal tubules were labeled with *Lotus tetragolonobus* lectin (LTL) (green), and nuclei were counterstained with DAPI (blue). Confocal images revealed the presence of cysts of variable sizes, with some cysts showing red or green staining, confirming their origin from the distal and proximal tubules, respectively. Notably, additional cysts were observed that were negative for both LTL and DBA markers (indicated by asterisks), suggesting that these cysts originate from other nephron segments beyond the proximal and distal tubules. No cysts were observed in littermates *Pkd2^+/fl^*. Scale bars = 200μm**. B)** *Left,* example confocal images of renal tubular epithelium sections harvest from *Pkd2^D509V/fl^* mice either treated with doxycycline (doxy.) for ten days or left untreated (control). Immunostaining performed using the primary cilia label anti-ADP-ribosylation factor-like 13B (ARL13B, polyclonal protientech 17711-1-AP) and the principal cell marker anti-aquaporin 2 (AQP2, monoclonal Abcam EPR21080). *Right,* analysis of the primary cilia density and length. Parenthesis indicate N per treatment group. *P* values result from unpaired two-tailed Welch’s t-tests.

## DISCUSSION

ADPKD is a channelopathy and ciliopathy— where both ion channel dysfunction and associated ciliary defects drive disease progression^24,47,67,68^. In this report, we assessed mechanistic impact and cystogenic propensity of the PKD2 D511V germline variant found in the patient population^47,69–71^ and our results clearly support the pathogenic classification of the mutation (https://pkdb.mayo.edu/)^36^. Our *in vitro* biochemical and cryo-EM structural analysis clearly indicates that D511V disrupts VSD electrostatic interactions which normally stabilize protein folding and channel assembly. The 3.1 Å cryo-EM structure of PKD2 D511V, together with its deposited atomic coordinates in the Protein Data Bank, provides a structural framework for developing protein stabilizing “corrector” molecules. In cultured cells and in animal models, the variant-induced folding defects abolish channel trafficking to the primary cilia, where the loss of channel function destabilizes the organelle structure, as supported by super resolution imaging and cilia electrophysiology results. D511V channels failed to function even after forcible reconstitution into synthetic lipid vesicles, which is consistent with previous analysis of this mutation in ER-localized channel populations^63,65^. When assessed in mice expressing the analogous mutation, we find severe hypomorphic PKD2 protein expression impairs primary cilia formation, indicating the destabilizing missense variant hinders channel stasis which precipitates organelle degeneration through regulatory pathways required for its maintenance. The following discussion points explore the implications of these findings in relation to mutation-induced molecular pathogenesis, and potential therapeutic strategies for treating ADPKD

### VSD variant impact on channel structural stability, trafficking and primary cilia maintenance

Our cryo-EM structural analysis suggests the D511V variant disrupts gating charge electrostatic interaction that are highly conserved among members of the superfamily of voltage-gated ion channels^59,72^. Neutralization of these interactions in Na_v_ and Ca_v_ channels are implicated in several other channelopathy diseases, including hypokalemic periodic paralysis, inherited pain syndromes forms of epilepsy and cardiac arrhythmias^59,73,74^. While several of VSD mutations alter the functional gating properties (opening, closing) of Ca_v_ and Na_v_ channels^75–77^, their role in stabilizing Kv channel folding and assembly is also established^78–81^. Our biochemical results suggest the dysregulation mechanism of the D511V variant falls within the latter category, causing channel protein destabilization. However, our CFE-GUV results also demonstrate that the D511V variant is incapable of channeling ions, even when artificially inserted in membranes. The D511V variant abolished PKD2 trafficking to primary cilia and disrupted ciliary morphology when expressed in cell lines, developing renal tissue, and adult kidney tubules. This broad ciliary degeneration phenotype reinforces the central role of primary cilia dysfunction in disease pathology. While several ADPKD-associated frame shift and truncating mutations are located in the PKD2 VSD, only one other missense variant has been implicated with the disease (**Supplemental Figure 1B**). Like D511V, L517R is also in the S3 transmembrane helix but does not directly participate in electrostatic interactions. Recent work has implicated defective L517R-cholesterol interactions, which results in channel exclusion from the primary cilia^22^. While these results present a compelling case for lipid-directed channel trafficking, the impact of the L517R variant on protein stability and *in vivo* protein expression was not assessed. Thus, future work should determine other ADPKD-associated variants also share the mechanism of cilia channel loss protein stabilization.

### Clinically relevant variant mouse model and insights for ADPKD therapeutic development

ADPKD is has dominant inheritance but is recessive at the cellular level, as cystogenesis occurs after renal cells acquire a second somatic mutation impacting either PKD1 or PKD2 alleles^82^. After identifying the significant destabilizing effect of the D511V variant *in vitro*, we hypothesize the mouse equivalent mutation (D509V) carry a significant phenotype. We observed that homozygous *Pkd2 ^D509V/D509V^* mice are embryonically lethal. This phenotype aligns with the phenotype observed in homozygous *Pkd2* null mice (*Pkd2^-/-^*) and from mice harboring mutations (*Pkd2^lrm4^ or Pkd2^L511R^*) that disrupt the normal PKD2 cilia trafficking^17,22,83–85^. Beyond the kidney, PKD2 channels function in sensory cilia of the embryonic node, which are essential for establishing asymmetric gene expression required for cardiac development and left-right patterning^17,83–85^. Thus, homozygous PKD2 protein destabilization and nodal cilia degeneration likely results in embryonic lethality due to loss of channel- and cilia-mediated Ca^2+^ signaling. Biochemical analysis of the mouse embryonic fibroblasts indicate variant alleles significantly reduces PKD2 channel protein expression. We attempted to reinstate variant protein levels in vitro by lowering the culture temperature conditions and pharmacologically inhibiting lysosomal and proteasomal degradation, but neither approach proved effective. Based on these findings, development of D511V-targeted molecular chaperones (“correctors”) that reinstate the variant protein stability may present a viable therapeutic strategy for ADPKD. We have resolved the PKD2 D511V cryo-EM structure at 3.1Å and shared the atomic coordinates (www.PDB.org) which may serve as a template for designing corrector molecules. Clinical trials are in progress for small molecule that rescue defective protein folding of PKD1^86^. However, these prototypic drugs are untested in patients carrying germline PKD2 variants, like D511V. Gene-corrector or gene-replacement strategies which reinstate PKD2 hypomorphic expression may also present a rational approach for patients carrying the voltage sensor varriant^87^. While PKD1 and PKD2 gene knockout mice are available to study disease pathogenesis, these models are less useful for on-target therapeutic development, as they completely ablate expression of polycystin channel genes^88,89^. Thus, the developed *Pkd2^D509V/fl^*mice address a need for ADPKD models based on a human variant to evaluate polycystin-targeted drug efficacy *in vivo*.

### Limitations of this study

As discussed in the introduction, subunits of PKD2 can form homomeric voltage-gated and Ca^2+^ modulated ion channels in the primary cilia^24^. Because of their tentability to functional analysis, we directed our experiments and interpreted our results primarily within the context of the homomeric PKD2 channels. Nonetheless, since PKD2 expression is integral to the formation of the PKD1-PKD2 channel complexes, we hypothesis that D511V hypomorphic expression and structural destabilizing effects would likely impair heteromeric PKD1-PKD2 complex assembly. Future studies examining the effects on PKD1-PKD2 channels will benefit from our established animal models expressing a clinically relevant ADPKD-causing PKD2 variant.

## Disclosure Statement and data availability statement

Data reported in this paper is deposited at the ARCH NU library and will be available without restriction upon acceptance of this manuscript (https://doi.org/XXXXX). The atomic coordinates for the PKD2 D511V variant channel have been deposited in the protein databank under accension codes PDB ID: 9NJS and EMDB ID EMD-49489 (*released upon publication)*. PDB validation reports are attached as supplement to this submission.

## Acknowledgements

We thank all laboratory personnel for their useful comments during the progress of this study. Q.W. is the recipient of the Jared J. Grantham Research Fellowship (American Society of Nephrology). P.G.D. was supported by the National Institute of Diabetes and Digestive and Kidney Diseases (R01 DK131118-01). These studies utilized resources provided by the NIDDK-sponsored Polycystic Kidney Disease Research Resource Consortium (PKD RRC) (1U54DK126114). The authors wish to thank members of the Qian laboratory for their helpful advice and Dr. Mauban at the University of Maryland School of Medicine’s Confocal Microscopy Core, Baltimore, Maryland (S10 OD026698) for imaging guidance.

---

**Supplemental Figure 1.**
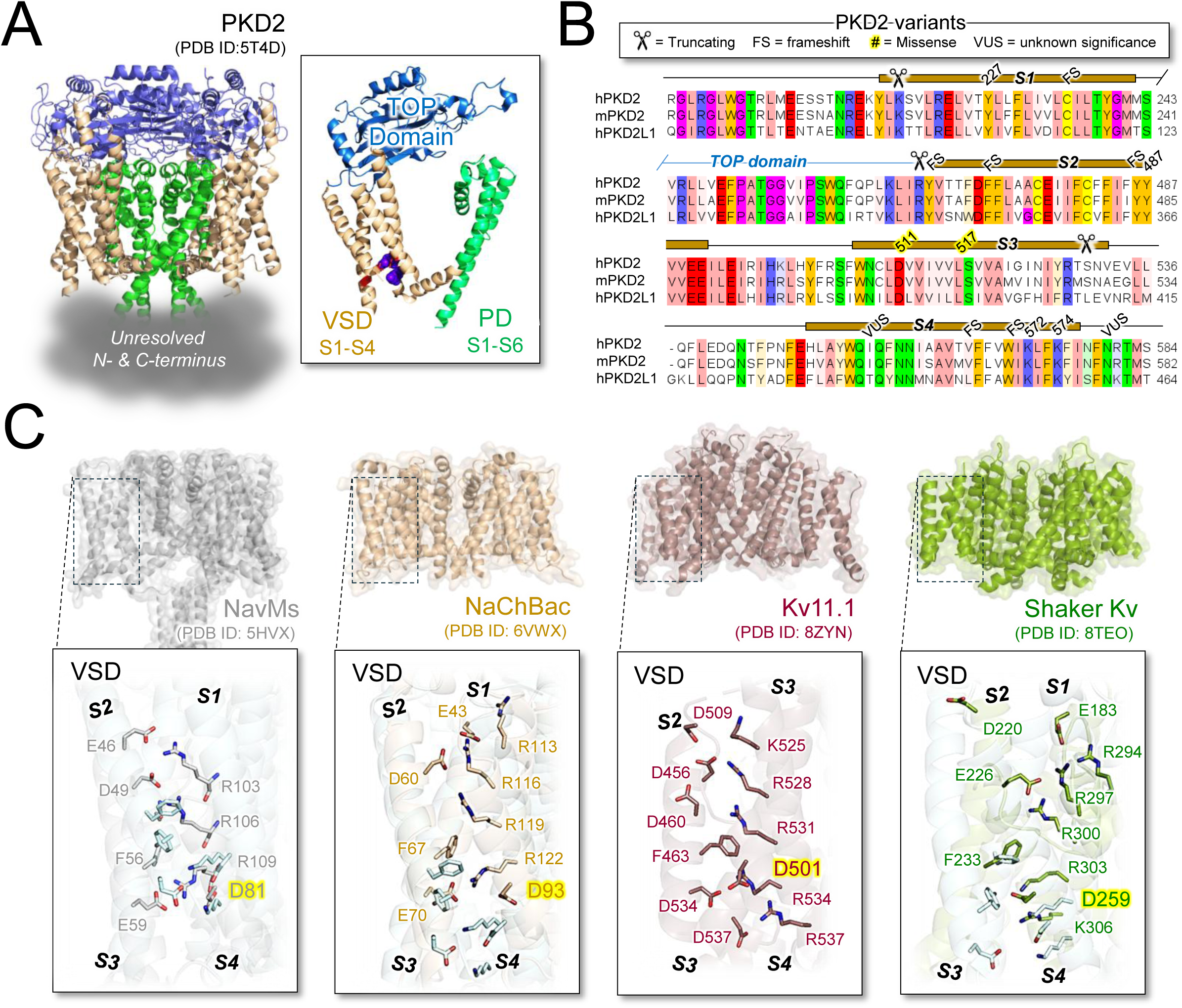
Conservation of gating charge electrostatic intactions within the PKD2 VSD and location of ADPKD-associated variants. **A**) Overview of the homotetrameric PKD2 channel structure highlighting the subunits domains. **B**) Amino acid aligment of polycystin (human PKD2, murine PKD2, human PKD2L1) voltage sensor domains generated using Clustal-omega and rendered in JalView. The barrels indicate transmembrane alpha helices found in the WT PKD2 cryo-EM structure and the location of ADPKD-assocated varriants are indicated^91^. Note the compleate TOP domain has been truncated from the aligment. **C**) Structural aligments of PKD2 with other homotetrameric voltage gated ion channels of divergen classes and species: NaChBac (Bacillus halodurans), NavMs (Magnetococcus marinus), Kv11.1 (human), and Shaker Kv (Drosophila melanogaster orthologous of Kv1.1)^92–95^.

**Supplemental Figure 2.**
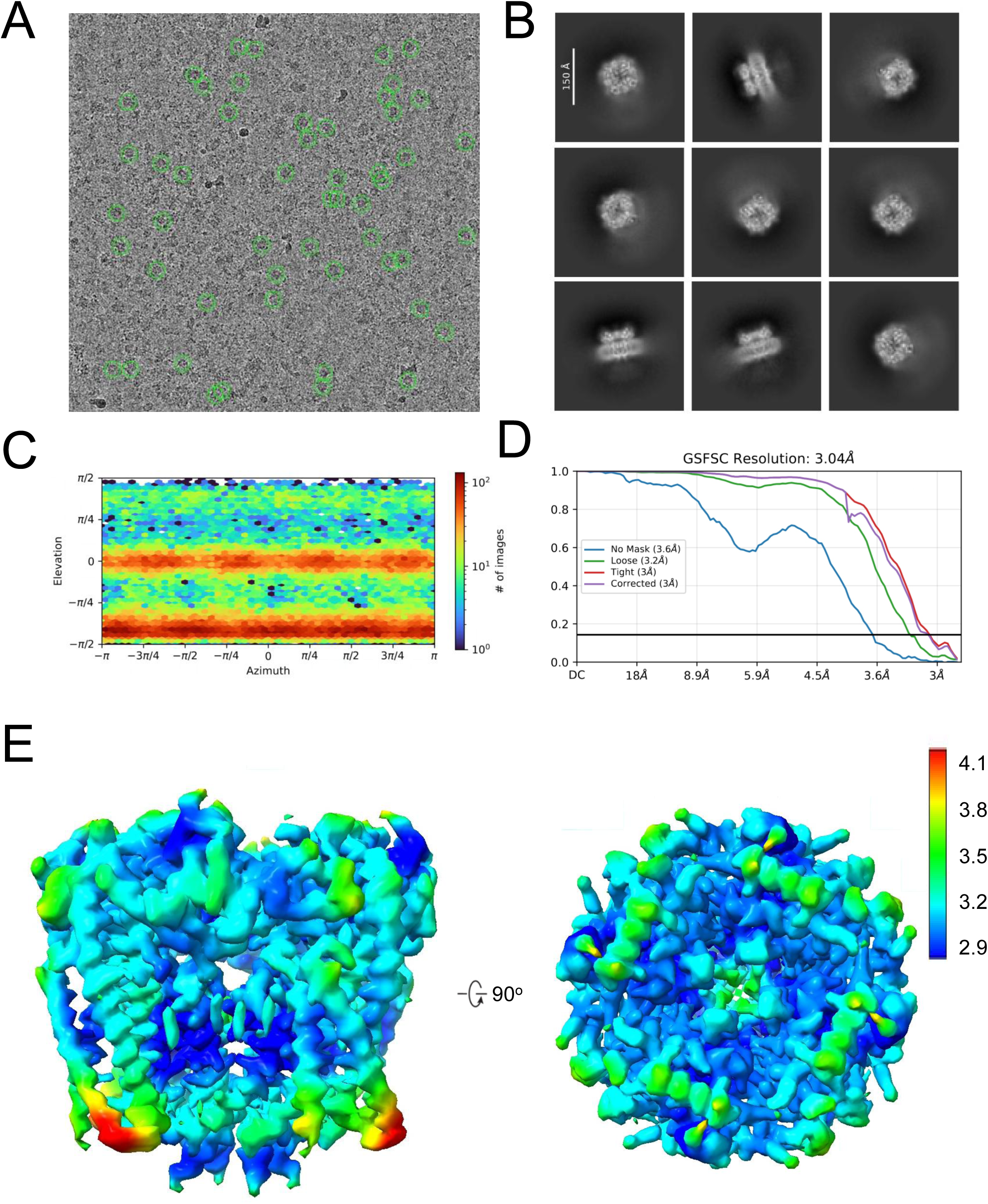
Cryo-EM data processing workflows used to determine the PKD2 D511V variant structure. **A**) representative raw micrograph shows polycystin-2 D511V channel particles. **B**) 2D classes of the D511V variant by Cryosparc show a sub-population of particles preserve the overall tetrameric architecture. However, fussy densities around the detergent belt and intracellular region are observable. **C**) Particle orientation plot by Cryosparc. **D**) Gold-standard FSC curves for resolution estimation and model validation. **E**) Local resolution plot. The local resolution in VSD, closing to V511, is the worst in the PKD2 tetramer while the channel pore reaches sub-3-Å resolution, suggesting destabilized VSD due to mutation effect.

**Supplemental Figure 3.**
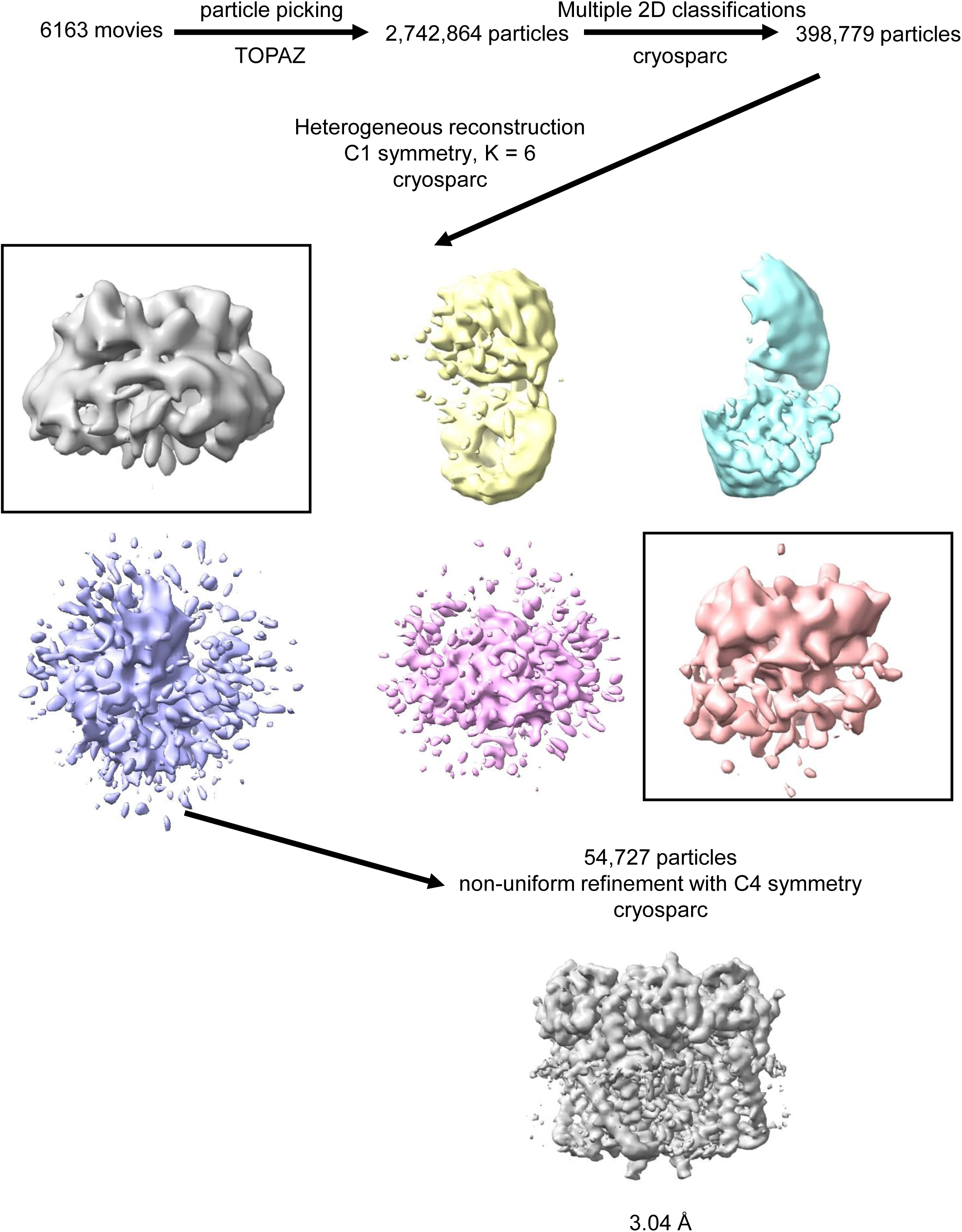
Reconstruction of human PKD2 D511V variant. Flow chart of image processing for the polycystin-2 D511V channel.

**Supplemental Figure 4.**
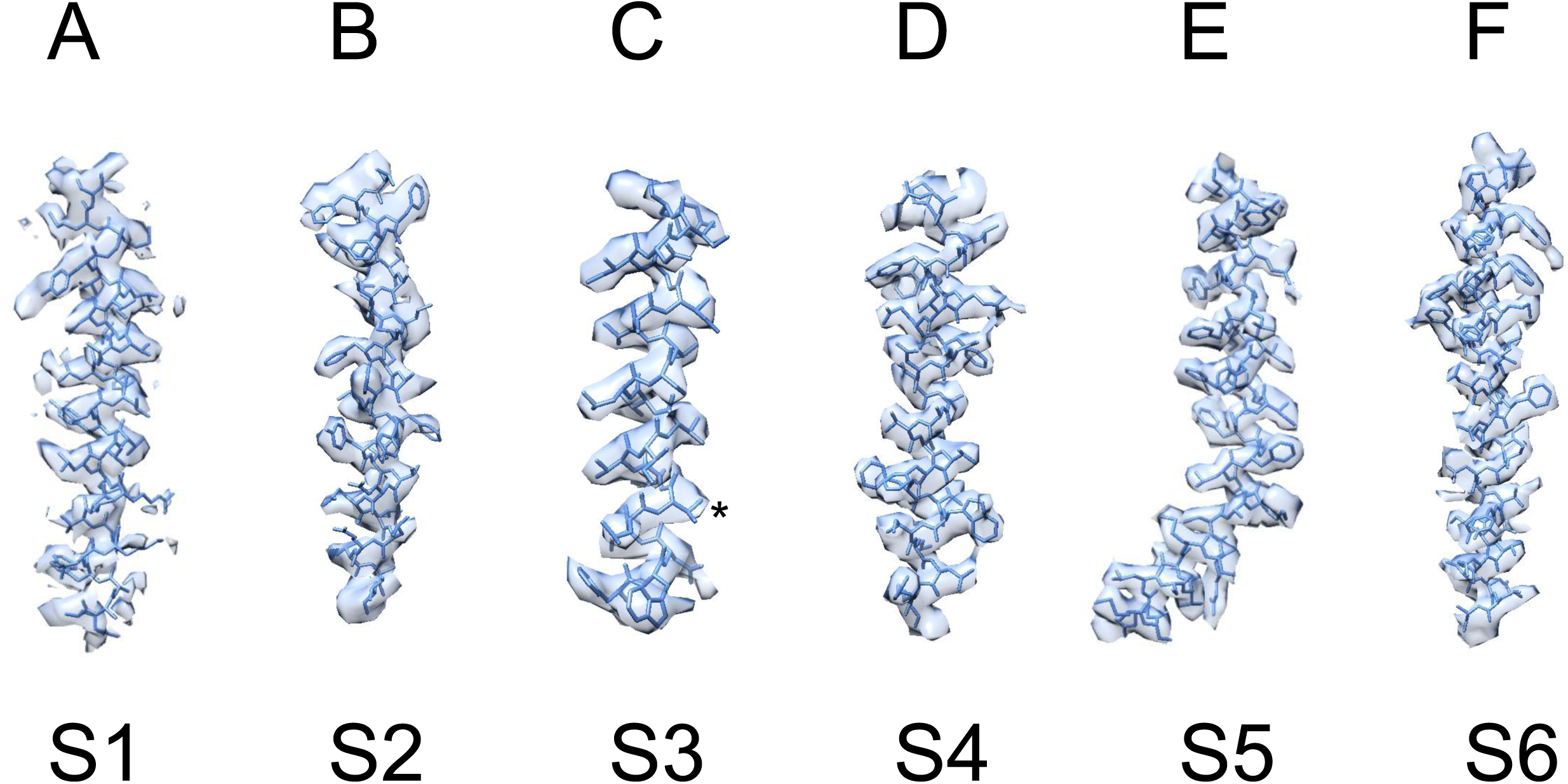
EM density maps of transmembrane regions from the PKD2 D511V variant. The maps are sharpened with a b factor of -100 Å^2^. The position of D511V is marked by asterisk.

**Supplemental Figure 5.**
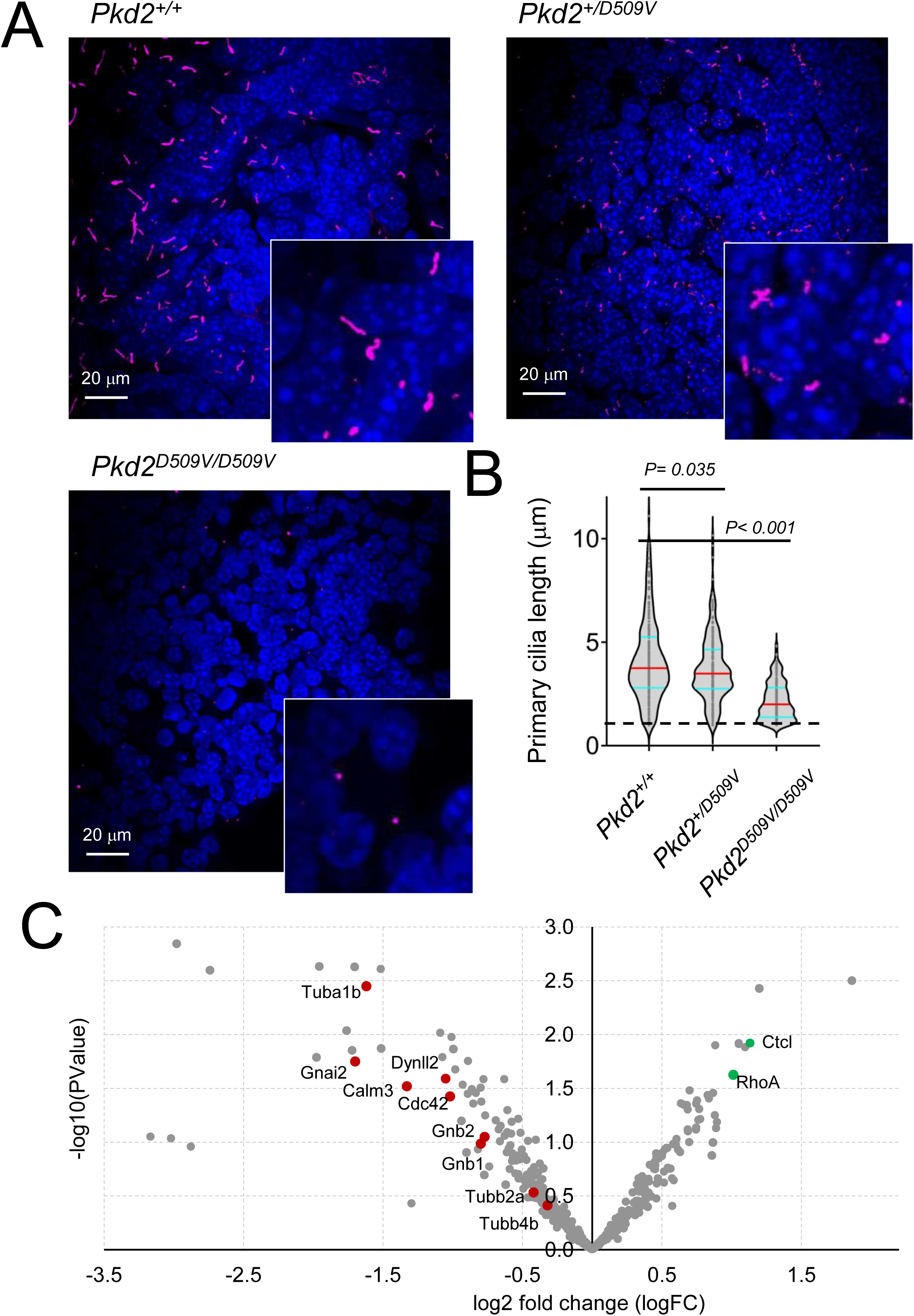
Attenuated renal cilia length in mice expressing the VSD variant. **A)** Immunohistochemistry confocal images of embryonic kidney (E14-E15) sections from mice expressing wild-type *Pkd2^+/+^* and the homozygous *Pkd2^D509V/D509V^*variant allele. Tissues were parfomaldyde fixed, cell nuclei were stained with DAPI and primary cilia were labeled with anti-ARL13B antibody (Protientech, 17711-1-AP). **B**) Violin plots of renal primary cilia length. Horizontal red line indicates the average and cyan lines indicate the 25 and 75 percentiles of each data set. P-values resulting from a one-way ANOVA statistical analysis are shown above the plots. Averages are shown at horozontal lines and error bars are equal to standard deviation. Dashed line indicates the ARL13B signal 1 μm minimum cilia length threshold. Parentheses indicate N per treatment group. P values result from unpaired two-tailed Welch’s t-tests. **C**) Volcano plot showing transcript-level differential expression between WT and homozygous PKD2-D509V E16.5 mouse kidneys. Each point represents an individual gene plotted by log₂ fold change on the x-axis and –log₁₀ p-value on the y-axis. Genes of interest linked to cytoskeletal regulation, centrosome/cilia stability, tubulin dynamics, and G-protein signaling are highlighted in red (Cdc42, Gnai2, Gnb1, Gnb2, Tubb4b, Tubb2a, Calm3, Dynll2, Tuba1b) or green (RhoA, Cltc), with all other genes shown in grey.

**Supplemental Figure 6.**
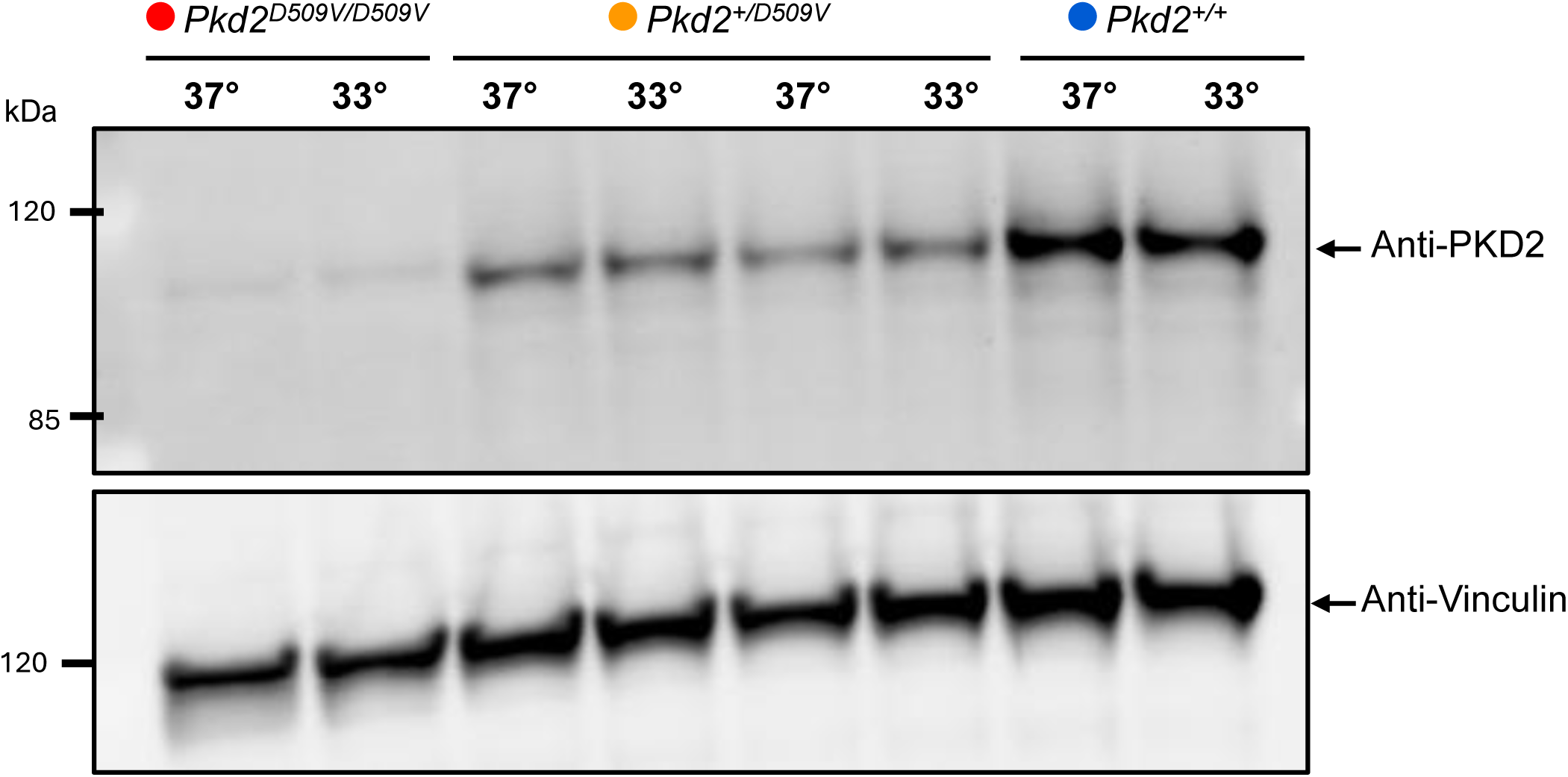
Polycystin variant protein expression under variable temperture. Western blot analysis of Pkd2 protein expression from MEF cells isolated from homozygous (red), heterozygous (orange) and wild-type (blue) embryos cultured at 37°C and 33°C (to rescue thermodynamic instability) in a 5% CO₂ incubator for 48 hours. Vinculin was used as a loading control. Notably, the expression levels of the Pkd2 D509V variant remain unchanged after culturing cells at 33°C, suggesting that the *Pkd2^D509V^* variant is not a temperature-sensitive folding mutant.

**Supplemental Figure 7.**
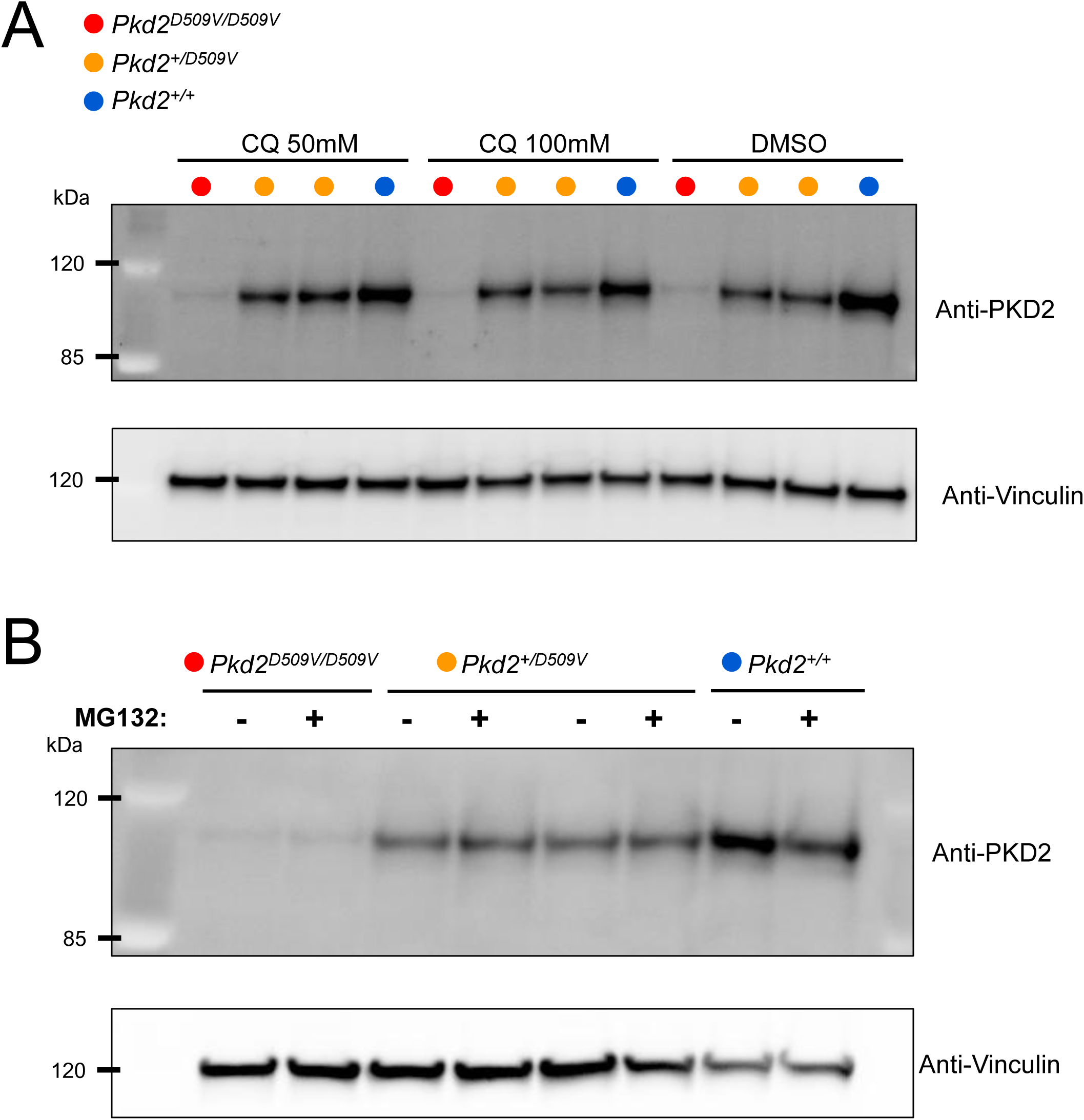
Polycystin variant protein expression after lysosomal or proteasomal inhibition. **A**) Immunoblot of Pkd2 protein from MEFs isolated from mice expressing the variant alleles after 24-hour treatment with DMSO (0.1% vehicle control) or lysosomal-degradation inhibitor chloroquine (CQ). Vinculin was used as loading control. **B**) As shown in A), immunoblots of MEFs after 6-hour treatmnet with DMSO or the proteasomal degradation-inhibitor, MG132.

**Supplemental Table 1.** Statistics of cryo-EM data acquisition and processing and model building.

| Data collection and processing of Human PKD2 D511V |  |
| --- | --- |
| Microscope | Titan Krios/K2 camera |
| Voltage (kV) | 300 |
| Defocus range ($\mu\text{m}$) | -0.6 – -3.6 |
| Pixel size ($\text{\AA}$) | 1.38 |
| Total electron dose ($\text{e}^-/\text{\AA}^2$) | 59 |
| Exposure time (s) | 13 |
| Number of movies | 6,163 |
| Number of frames per movie | 40 |
| Initial particle number | 2,742,864 |
| Final particle number | 54,727 |
| Resolution (unmasked, $\text{\AA}$) | 3.6 |
| Resolution (masked, $\text{\AA}$) | 3.1 |
| Number of atoms | 16,676 |
| R.M.S deviation |  |
| Bond length ($\text{\AA}$) | 0.006 |
| Bond angles ($^\circ$) | 0.729 |
| Ramachandran |  |
| Favored (%) | 94.48% |
| Allowed (%) | 5.52% |
| Outlier (%) | 0.00% |
| Molprobity score | 2.07 |

**Supplemental Table 2.** Results from time pregnancies at E14.5 and P0 resulting from crossing heterozygous *Pkd2^+/D509V^*mice. We observed that heterozygous *Pkd2^+/D509V^* mice appear normal and fertile (at ∼ 6 months of age); however, no viable *Pkd2 ^D509V/D509V^* homozygous pups were born after the interbreeding of heterozygous mice. Timed pregnancy studies showed that Mendelian ratios were partially restored when embryos were harvested at E14.5.

| Number of viable pups at P0 |  |  |  |
| --- | --- | --- | --- |
|  | *Pkd2*<sup>+/+</sup><br>Wild-type | *Pkd2*<sup>+/D509V</sup><br>Heterozygous | *Pkd2*<sup>D509V/D509V</sup><br>Homozygous |
| **Observed** | **13 (46.5%)** | **14 (50%)** | **0 (0%)** |
| *Expected* | 7 (25%) | 14 (50%) | 7 (25%) |
| Number of viable embryos at E14.5 |  |  |  |
| **Observed** | **15 (23%)** | **38 (57%)** | **13 (20%)** |
| *Expected* | ~17 (25%) | ~32 (50%) | ~17 (25%) |

